# Genome-wide local ancestry and the functional consequences of admixture in African and European cattle populations

**DOI:** 10.1101/2024.06.20.599852

**Authors:** Gillian P. McHugo, James A. Ward, Said Ismael Ng’ang’a, Laurent A.F. Frantz, Michael Salter-Townshend, Emmeline W. Hill, Grace M. O’Gorman, Kieran G. Meade, Thomas J. Hall, David E. MacHugh

**Author notes:** **Address for correspondence**: D. E. MacHugh, Animal Genomics Laboratory, UCD School of Agriculture and Food Science, University College Dublin, Belfield, Dublin, D04 V1W8, Ireland.

## Abstract

*Bos taurus* (taurine) and *Bos indicus* (indicine) cattle diverged at least 150,000 years ago and, since that time, substantial genomic differences have evolved between the two lineages. During the last two millennia, genetic exchange in Africa has resulted in a complex tapestry of taurine-indicine ancestry, with most cattle populations exhibiting varying levels of admixture. Similarly, there are several Southern European cattle populations that also show evidence for historical gene flow from indicine cattle, the highest levels of which are found in the Central Italian White breeds. Here we use two different software tools (MOSAIC and ELAI) for local ancestry inference (LAI) with genome-wide high-and low-density SNP array data sets in hybrid African and Italian cattle populations and obtained broadly similar results despite critical differences in the two LAI methodologies used. Our analyses identified genomic regions with elevated levels of retained or introgressed ancestry from the African taurine, European taurine, Asian indicine lineages. Functional enrichment of genes underlying these ancestry peaks highlighted biological processes relating to immunobiology and olfaction, some of which may relate to differing susceptibilities to infectious diseases, including bovine tuberculosis, East Coast fever, and tropical theileriosis. Notably, for retained African taurine ancestry in admixed trypanotolerant cattle we observed enrichment of genes associated with haemoglobin and oxygen transport. This may reflect positive selection of genomic variants that enhance control of severe anaemia, a debilitating feature of trypanosomiasis disease, which severely constrains cattle agriculture across much of sub-Saharan Africa.

## Introduction

Long acknowledged in plants (Anderson, 1949), gene flow and hybridisation between interfertile taxa are increasingly recognised as important evolutionary processes in animals (Hedrick, 2013; Payseur and Rieseberg, 2016; Taylor and Larson, 2019). Genetic exchange between populations can provide an abundant source of new functional genomic variation—both adaptive and maladaptive—that can generate novel combinations of alleles at individual genes, and interacting gene loci, thereby altering gene regulatory networks, biochemical pathways, physiological outputs, and ultimately phenotypic outcomes (Arnold and Kunte, 2017; Edelman and Mallet, 2021; Tigano and Friesen, 2016). In this respect, hybrid zones, where evolutionary distinct but interfertile animal taxa interact to produce admixed populations, represent natural laboratories for evolutionary studies (Hewitt, 1988). It has also been observed that gene flow, reticulate evolution, and admixture between distinct lineages and from wild congeners are common features of many domestic animal species, including pigs (*Sus scrofa*), dogs (*Canis familiaris*), sheep (*Ovis aries*), goats (*Capra hircus*), and chickens (*Gallus gallus*) (Frantz *et al*, 2015; Freedman and Wayne, 2017; Lv *et al*, 2022; Pogorevc *et al*, 2024; Wang *et al*, 2020b).

Cattle were domesticated from the now extinct aurochs (*Bos primigenius*) (Bailey *et al*, 1996) and humpless *Bos taurus* (taurine) cattle were some of the first large ruminants to be domesticated 10−11,000 years ago in the Fertile Crescent region (Conolly *et al*, 2011; Larson *et al*, 2014; Zeder, 2017). Approximately 2,000 years later, humped *Bos indicus* (indicine or zebu) cattle were domesticated in present-day Pakistan (Utsunomiya *et al*, 2019) and analyses of genome-scale DNA sequence data show that the *B. taurus* and *B. indicus* lineages likely diverged 150–500 kya (Chen *et al*, 2018; Wang *et al*, 2018; Wu *et al*, 2018). Consequently substantial genomic differences have evolved between the two subspecies, making hybrid cattle an excellent resource for addressing fundamental scientific questions concerning the role of gene flow, admixture, and introgression in mammalian microevolution (Bahbahani *et al*, 2017; Chen *et al*, 2018; Chen *et al*, 2023; Flori *et al*, 2014; Friedrich *et al*, 2023; Kim *et al*, 2020; Kwon *et al*, 2022; Mbole-Kariuki *et al*, 2014; McTavish and Hillis, 2014; Ward *et al*, 2022; Wu *et al*, 2018).

Cattle populations from several regions around the globe exhibit evidence of *B. taurus*/*B. indicus* admixture, although gene flow and genomic introgression between the two subspecies is most well understood and surveyed in Africa (Decker *et al*, 2014; Hanotte *et al*, 2002; Hanotte *et al*, 2000; Kim *et al*, 2020; MacHugh *et al*, 1997). Domestication and the subsequent spread and interactions of different taurine and indicine cattle populations has resulted in gradients of *B. taurus* and *B. indicus* ancestry across the African continent (Hanotte *et al*, 2002; Mwai *et al*, 2015). There are approximately 150 breeds of indigenous cattle in sub-Saharan Africa and African cattle represent a complex tapestry of African *B. taurus* and Asian *B. indicus* ancestry, with some populations also exhibiting significant non-African *B. taurus* genetic influence (Hanotte *et al*, 2002; Kim *et al*, 2020; MacHugh *et al*, 1997). The indigenous *B. taurus* cattle of Africa are generally adapted to humid and subhumid zones associated with sedentary subsistence farming and face particular disease challenges as a consequence (FAO, 2015). As a result of their longer history of exposure and adaptation to high pathogen and parasite burdens on the continent, African *B. taurus* cattle have several advantages over *B. indicus* cattle in terms of disease tolerance and resistance (de Clare Bronsvoort *et al*, 2013; Mwai *et al*, 2015). Cattle that are predominantly indicine in ancestry are normally transhumant livestock adapted to the arid and semi-arid regions of the continent and are favoured by many farmers due to their larger size and higher production yields, while hybrid populations tend to inhabit environments somewhere between these extremes (FAO, 2015; Mwai *et al*, 2015).

One particularly important disease for African cattle is African animal trypanosomiasis (AAT) or nagana, a wasting disease caused by parasitic protozoa of the genus *Trypanosoma* transmitted by biting insect vectors such as tsetse flies (*Glossina* spp.), which causes fever, severe weight loss and anaemia (MacGregor *et al*, 2021; Steverding, 2008). Cattle agriculture in sub-Saharan Africa is severely constrained by AAT because, even with the availability of trypanocidal drugs, the high susceptibility of many breeds to trypanosomiasis renders them unproductive in regions with significant tsetse burdens (Berthier *et al*, 2015; Yaro *et al*, 2016). However, some African *B. taurus* breeds have a tolerance of trypanosome infection termed “trypanotolerance”, which enables these cattle to control parasitaemia and anaemia, making them more productive than trypanosusceptible breeds in many areas of West and Central Africa (Berthier *et al*, 2015; Murray and Black, 1985). These trypanotolerant populations, which include the longhorn N’Dama and shorthorn Baoule, Lagune, and Somba breeds, are therefore an important genetic resource as they are uniquely suited to livestock production in these areas (Berthier *et al*, 2015; Yaro *et al*, 2016).

Trypanotolerance has been shown to be a heritable multigenic trait, with variability in tolerance among individual animals within trypanotolerant populations (Kambal *et al*, 2023). Some African *B. taurus*/*B. indicus* hybrid cattle breeds are also known to exhibit trypanotolerance; however, trypanotolerant breeds with high levels of *B. taurus* ancestry have a greater capacity to control anaemia, while hybrid animals exhibit intermediate levels of control compared to trypanosusceptible *B. indicus* breeds (Bahbahani *et al*, 2018; Berthier *et al*, 2015). The genomic architecture of trypanotolerance in cattle remains poorly understood, although some candidate genes have been proposed, and identification of genes and genomic regulatory elements (GREs) underpinning the trait may facilitate introduction or enhancement of the trait via genome-enabled breeding or genome editing (Yaro *et al*, 2016).

In contrast to the complex nature of African cattle ancestry, the majority of European cattle populations are comprised of pure European *B. taurus* ancestry; however, there are several breeds in Southern Europe that are known to exhibit modest levels of African *B. taurus* and/or *B. indicus* ancestry (Upadhyay *et al*, 2019). The most well-characterised of these, which also have the highest levels of indicine admixture, are the group of populations known as Central Italian White cattle (Barbato *et al*, 2020). Compared to temperate taurine cattle, *B. indicus* cattle have enhanced heat and drought tolerance and introgression of genomic variants from *B. indicus* into Central Italian White cattle may have made these breeds better adapted to extreme summer heat events on the Italian peninsula (Hooyberghs *et al*, 2019).

Admixture and introgression among populations can be studied at a sub-chromosomal level using statistical methods for surveying locus-specific or local ancestry, which in contrast to global ancestry proportions, corresponds to the ancestry of specific genomic segments that consist of unbroken ancestry blocks from different donor populations (Gompert and Buerkle, 2013). A range of methods for local ancestry inference (LAI) using genome-scale data have been developed (Tan and Atkinson, 2023; Wu *et al*, 2021). Two widely used software tools are *Efficient Local Ancestry Inference* (*ELAI*) (Guan, 2014) and *MOSAIC Organizes Segments of Ancestry In Chromosomes* (*MOSAIC*) (Salter-Townshend and Myers, 2019). ELAI fits a two-layer hidden Markov model (HMM) that allows ancestry switching anywhere along the genome; however, it requires the donor reference populations and the approximate number of generations since the admixture occurred to be preassigned. Additionally, the donor reference populations should be as genetically similar to the original source populations as possible. MOSAIC also fits a two-layer HMM but employs a different strategy that determines how closely related each segment of chromosome in every admixed individual genome is to chromosomal segments in individual genomes from potential donor reference populations and infers a stochastic relationship between donor reference panels and mixing populations. Unlike other methods, MOSAIC does not require the donor reference populations to be direct surrogates for the original source populations and it can also infer the number of generations since the start of an admixture process. However, the MOSAIC algorithm requires phased haplotypes and a recombination rate map.

For the present study we performed a range of population genomics analyses and comparative LAI using the ELAI and MOSAIC software tools with a panel of African and European cattle breeds that exhibit varying levels of African taurine, European taurine, and Asian indicine ancestries. Two different genome-wide SNP data sets were used: a high-density SNP data set consisting of more than 600,000 SNPs and a low-density data set encompassing approximately 30,000 SNPs. These analyses allowed us to assess the ELAI and MOSAIC algorithms as tools for LAI in admixed cattle. We were also able to systematically catalogue and functionally evaluate genomic regions exhibiting evidence for elevated levels of introgressed or retained ancestry from the three cattle lineages.

## Materials and Methods

### High-density genome-wide cattle SNP data sets

For this study new Illumina^®^ BovineHD 777K BeadChip SNP data sets were generated for 39 African cattle (23 Somba, 8 N’Dama and 8 Boran). The Somba breed data were obtained using DNA samples previously published as part of a microsatellite-based survey of cattle genetic diversity (Freeman *et al*, 2004) and were generated by Weatherbys Scientific (Naas, Ireland) using standard procedures for Illumina^®^ SNP array genotyping. The N’Dama and Boran data were obtained using cattle DNA samples from a trypanosome challenge time-course experiment (O’Gorman *et al*, 2009) and were generated by Neogen Europe (Ayr, Scotland) also using standard procedures. Additional Illumina^®^ BovineHD 777K BeadChip data sets were obtained from published studies (Bahbahani *et al*, 2017; Barbato *et al*, 2020; Upadhyay *et al*, 2017; Verdugo *et al*, 2019; Ward *et al*, 2022; Wragg *et al*, 2022) and the Web-Interfaced next generation Database Exploration (WIDDE) repository Sempéré *et al* (2015).

The total data set consisted of high-density 777K SNP data for 1,030 cattle before filtering and 24 different populations were represented, including three European *B. taurus* populations (Holstein Friesian, Angus, and Jersey); three African *B. taurus* populations (Muturu, Lagune, and Guinean N’Dama); three *B. indicus* populations (Tharparkar, Gir, and Nelore); five European hybrid populations (Romagnola, Chianina, Marchigiana, Maremmana, and Alentejana); five trypanotolerant African hybrid populations (hybrid N’Dama, Borgou, Somba, Keteku, and Sheko) and five trypanosusceptible African hybrid populations (Ankole, Nganda, East African Shorthorn Zebu, Karamojong, and Boran). The cattle BovineHD 777K SNP data were converted to binary PLINK files with Illumina^®^ allele coding for the FORWARD strand as required using PLINK (v. 1.90 beta 6.25) (Chang *et al*, 2015) and SNPchiMp (v. 3) (Nicolazzi *et al*, 2015). The sample data were then merged with PLINK (v. 1.90 beta 6.25).

Figure 1 illustrates the overall study workflow including the genome assembly updating, data preparation, and filtering steps, which are described in the following subsections and that were implemented prior to the population genomics analyses. Table 1 shows the taxonomic, breed, geographical, sample number (pre-and post-SNP data filtering), and sources for the BovineHD 777K BeadChip SNP data sets. There was a total of 750 individual animal BovineHD 777K BeadChip SNP data sets retained after filtering.

**Figure 1.**
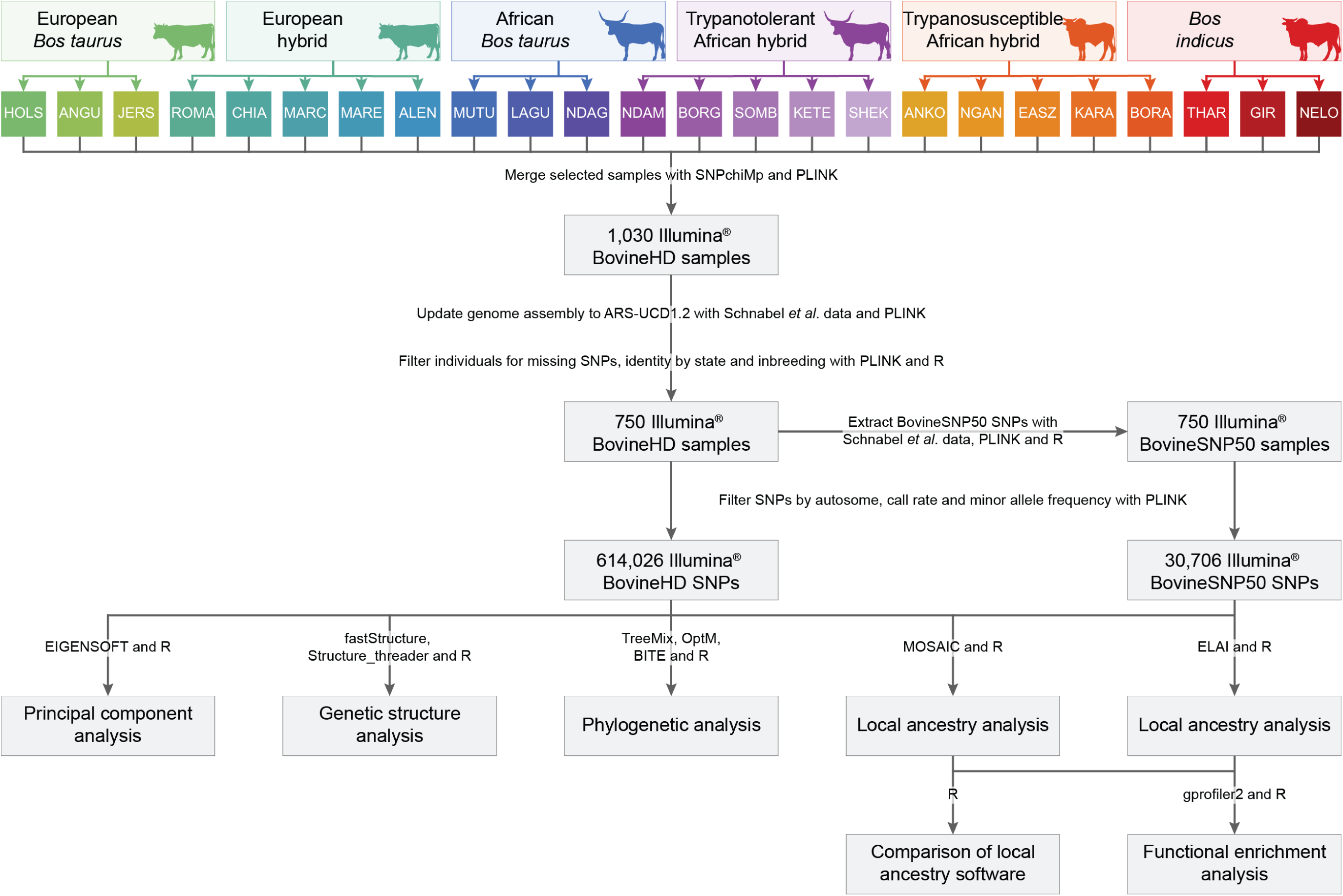
Diagram showing study workflow. Cattle images by Tracy A. Heath, T. Michael Keesey and Steven Traver via phylopic.org. Colours were generated from the khroma R package (v. 1.10.0) (Frerebeau, 2023).

**Table 1.** Group, code, population, origin, number of samples pre-and post-filtering and sources of SNP data used in this study.

| Group | Code | Population | Country of origin | No. pre-filtering | No. post-filtering | Source* |
| --- | --- | --- | --- | --- | --- | --- |
| European<br>*Bos taurus* | HOLS | Holstein Friesian | Netherlands | 60 | 36 | a |
| European<br>*Bos taurus* | ANGU | Angus | United Kingdom | 42 | 26 | a |
| European<br>*Bos taurus* | JERS | Jersey | United Kingdom | 38 | 23 | a |
| European hybrid | ROMA | Romagnola | Italy | 51 | 51 | b, a |
| European hybrid | CHIA | Chianina | Italy | 19 | 19 | b, c |
| European hybrid | MARC | Marchigiana | Italy | 13 | 13 | b |
| European hybrid | MARE | Maremmiana | Italy | 9 | 5 | c |
| European hybrid | ALEN | Alentejana | Portugal | 11 | 6 | d, c |
| African<br>*Bos taurus* | MUTU | Muturu | Nigeria | 39 | 19 | e |
| African<br>*Bos taurus* | LAGU | Lagune | Benin | 5 | 5 | a |
| African<br>*Bos taurus* | NDAG | N'Dama Guinea | Guinea | 70 | 47 | e, a, f |
| Trypanotolerant<br>African hybrid | NDAM | N'Dama hybrid | Unspecified,<br>Togo, Kenya | 65 | 41 | b, d, g |
| Trypanotolerant<br>African hybrid | BORG | Borgou | Benin | 50 | 50 | e |
| Trypanotolerant<br>African hybrid | SOMB | Somba | Benin | 33 | 23 | g |
| Trypanotolerant<br>African hybrid | KETE | Keteku | Nigeria | 22 | 22 | e |
| Trypanotolerant<br>African hybrid | SHEK | Sheko | Ethiopia | 16 | 16 | a |
| Trypanosusceptible<br>African hybrid | ANKO | Ankole | Uganda | 50 | 25 | f |
| Trypanosusceptible<br>African hybrid | NGAN | Nganda | Uganda | 53 | 27 | f, e |
| Trypanosusceptible<br>African hybrid | EASZ | East African<br>Shorthorn Zebu | Kenya | 111 | 111 | f, e |
| Trypanosusceptible<br>African hybrid | KARA | Karamojong | Uganda | 16 | 16 | f |
| Trypanosusceptible<br>African hybrid | BORA | Boran | Kenya | 90 | 90 | h, g |
| *Bos indicus* | THAR | Tharparkar | Pakistan | 13 | 13 | b |
| *Bos indicus* | GIR | Gir | India | 85 | 28 | a, f |
| *Bos indicus* | NELO | Nelore | Brazil | 69 | 38 | a, c, f |
| **Total** |  |  |  | 1030 | 750 |  |

\* a Sempéré *et al* (2015), b Barbato *et al* (2020), c Upadhyay *et al* (2017), d Verdugo *et al* (2019), e Ward *et al* (2022), f Bahbahani *et al* (2017), g this study, h Wragg *et al* (2022).

### Updating the bovine genome assembly

The BovineHD 777K BeadChip SNP locations were updated from the UMD3.1 bovine genome assembly to the current assembly ARS-UCD1.2 (Rosen *et al*, 2020) using coordinates from the National Animal Genome Research Program (NAGRP) Data Repository *genotyping array SNP mapping to ARS-UCD1.2 resource* (Schnabel, 2018) and PLINK (v. 1.90 beta 6.25).

### Data preparation and filtering

#### Generation of a low-density SNP array data set

To produce a comparative low-density SNP array data set, the high density BovineHD 777K SNP data set was downsampled to the subset of the 46,713 SNPs in common with the Illumina^®^ Bovine SNP50 BeadChip using PLINK (v. 1.90 beta 6.25). A list of the Bovine SNP50 BeadChip SNPs from the NAGRP Data Repository (Schnabel, 2018) was used for this purpose and modified with dplyr (v. 1.1.2) (Wickham *et al*, 2023a) and readr (v. 2.1.4 (Wickham *et al*, 2023b) with R (v. 4.3.0) (R Core Team, 2023).

#### Missing SNP removal

Individual animals that had missing SNP call rates exceeding 0.95 from the low-density data set were removed using a missing genotype filter with PLINK (v. 1.90 beta 6.25). The same set of animals were also removed from the high-density data set.

#### Removal of duplicate samples by identity-by-state filtering

Duplicate samples present in two or more data sources were removed using PLINK (v. 1.90 beta 6.25) and a previously described identity-by-state methodology (Browett *et al*, 2018). The method was modified to select one from each pair of animals that had an identity-by-state value greater than or equal to 0.99 using dplyr (v. 1.1.2), readr (v. 2.1.4), and stringr (v. 1.5.0) (Wickham, 2023) with R (v. 4.3.0). The resulting list of sample duplicates were removed from the high-and low-density data sets with PLINK (v. 1.90 beta 6.25). The results were visualised using dplyr (v. 1.1.2), ggh4x (v. 0.2.4) (van den Brand, 2023), ggplot2 (v. 3.4.2) (Wickham, 2009), ggtext (v. 0.1.2) (Wilke and Wiernik, 2022), readr (v. 2.1.4), and stringr (v. 1.5.0) with R (v. 4.3.0). Colours were generated from viridis (v. 0.6.3) (Garnier *et al*, 2023) and khroma (v. 1.10.0).

#### Removal of admixed animals from the reference populations

An inbreeding analysis in PLINK (v. 1.90 beta 6.25) was used to remove animals that showed evidence for significant admixture in the reference populations (three European *B. taurus* populations: Holstein Friesian, Angus, and Jersey; three African *B. taurus* populations: Muturu, Lagune, and Guinean N’Dama; and three *B. indicus* populations: Tharparkar, Gir, and Nelore). To do this, outlier animals with statistically lower inbreeding values than the rest of the population were identified via boxplots using dplyr (v. 1.1.2), readr (v. 2.1.4), and tidyr (v. 1.3.0) (Wickham *et al*, 2023d) with R (v. 4.3.0). The resulting list of animals were removed from the high-and low-density data sets. A systematic inbreeding analysis was then performed with PLINK (v. 1.90 beta 6.25) and the output was modified using dplyr (v. 1.1.2) and readr (v. 2.1.4) with R (v. 4.3.0) to identify animals with the top 25 and bottom 25 inbreeding values across the three European *B. taurus* populations (Holstein Friesian, Angus, and Jersey). These samples were then removed from the high- and low-density data sets to balance the numbers of animals across the reference groups. A final inbreeding analysis of the low-density data set after the filters were applied was performed to compare the results with those of the high-density data set. Results from the inbreeding analyses were visualised using dplyr (v. 1.1.2), ggh4x (v. 0.2.4), ggplot2 (v. 3.4.2), ggtext (v. 0.1.2), and readr (v. 2.1.4), with R (v. 4.3.0). Colours were generated from viridis (v. 0.6.3) and khroma (v. 1.10.0).

#### Filtering of SNPs by call rate and minor allele frequency

The high and low-density data sets were filtered to retain autosomal SNPs with a minimum call rate of 95% and minor allele frequency (MAF) of at least 5% with PLINK (v. 1.90 beta 6.25). The methodologies used for this process have been described in a previous study (McHugo *et al*, 2019).

### Population genomics analyses

#### Principal component analysis

Principal component analysis (PCA) was performed for the high and low-density data sets using smartpca after file conversion with convertf, both part of EIGENSOFT package (v. 7.1.2) (Patterson *et al*, 2006). The results were visualised using dplyr (v. 1.1.2), ggplot2 (v. 3.4.2), patchwork (v. 1.1.2) (Pedersen, 2023), readr (v. 2.1.4), stringr (v. 1.5.0), and tidyr (v. 1.3.0) with R v. 4.3.0). Colours were generated from viridis (v. 0.6.3) and khroma (v. 1.10.0).

#### Genetic structure analysis

Genetic structure analysis was performed for the high and low-density data sets using structure_threader (v. 1.3.4) (Pina-Martins *et al*, 2017) with fastStructure (v. 1.0) (Raj *et al*, 2014). The structure analysis was carried out with the model complexity or number of populations (*K*) set from 2 to 25. The chooseK function was used to test the outputs to find a range of values of *K* that best accounted for the structure in the data (Raj *et al*, 2014). The results were visualised using dplyr (v. 1.1.2), ggh4x (v. 0.2.4), ggplot2 (v. 3.4.2), ggtext (v. 0.1.2), magick (v. 2.8.1) (Ooms, 2023) with ImageMagick (v. 6.9.12.96) (ImageMagick Studio LLC, 2023), magrittr (v. 2.0.3) (Bache and Wickham, 2022), patchwork (v. 1.1.2), readr (v. 2.1.4), stringr (v. 1.5.0) and tidyr (v. 1.3.0) with R (v. 4.3.0). Colours were generated from viridis (v. 0.6.3) and khroma (v. 1.10.0).

#### Phylogenetic analysis

An additional sample of three gaur (*B. gaurus*) that were genotyped using the BovineHD 777K BeadChip (Sempéré *et al*, 2015; Verdugo *et al*, 2019) were available to use as an outgroup. After the pre-processing steps described above were performed to convert the data to binary PLINK files with Illumina^®^ allele coding for the FORWARD strand on the ARS-UCD1.2 bovine genome assembly, the *B. gaurus* data set was filtered with PLINK (v. 1.90 beta 6.25) to retain only SNPs present in the high density data set. The gaur data were then merged with the high-density data set and an additional filter was applied with PLINK (v. 1.90 beta 6.25) to retain autosomal SNPs with a minimum call rate of 95% and minor allele frequency of at least 5%. A gzipped allele frequency cluster file was produced with PLINK (v. 1.90 beta 6.25) and the resulting file was converted to TreeMix format using the plink2treemix python script provided with the TreeMix software package (v. 1.13) (Pickrell and Pritchard, 2012).

Phylogenetic analysis was performed for both the high and low-density SNP data sets using TreeMix (v. 1.13) with the number of migration edges (*m*) set from 1 to 15 for ten iterations using windows of SNPs (*k*) increasing from 100 to 1000 by increments of 100. The OptM package (v. 0.1.6) (Fitak, 2021) was used with R (v. 4.3.0) to calculate the mean and standard deviation (SD) across the 10 iterations for the composite likelihood (*L*(*m*)), proportion of variance explained and the second-order rate of change (*Δm*) across migration edges (*m*). The results were visualised using dplyr (v. 1.1.2), ggplot2 (v. 3.4.2), ggtext (v. 0.1.2) and patchwork (v. 1.1.2) with R (v. 4.3.0). The BITE R package (v. 1.2.0008) (Milanesi *et al*, 2017) was also used to generate a Unix shell script customised to perform 100 TreeMix bootstrap replicates for the selected numbers of migration edges (*m*). The results were visualised using ape (v. 5.7.1) (Paradis and Schliep, 2019), dplyr (v. 1.1.2), ggh4x (v. 0.2.4), ggnewscale (v. 0.4.9) (Campitelli, 2023), ggplot2 (v. 3.4.2), ggtext (v. 0.1.2), ggtree (v. 3.9.0.1) (Yu, 2022), patchwork (v. 1.1.2), stringr (v. 1.5.0), tidytree (v. 0.4.4) (Yu, 2022), and treeio (v. 1.25.2) (Wang *et al*, 2020a) with R (v. 4.3.0) and a modified version of the script provided with TreeMix. Colours were generated from viridis (v. 0.6.3) and khroma (v. 1.10.0).

### Local ancestry estimation

#### MOSAIC analysis

The high and low-density SNP data sets were separated by chromosome with PLINK (v. 1.90 beta 6.25) and each chromosome was phased with SHAPEIT (v. 2.r904) (O’Connell *et al*, 2014). The resulting segregated chromosome SNP data files were converted to MOSAIC format using the R script provided with MOSAIC (v. 1.5.0) (Salter-Townshend and Myers, 2019) and R (v. 4.3.0). Recombination rate files were prepared from a cattle recombination map (Ma *et al*, 2015) using an R script adapted from Ward *et al* (2022) with R (v. 4.3.0). For each hybrid population three-way local ancestry analysis was performed across all autosomes without *F*_ST_ estimation and assuming an effective population size (*N_e_*) of 400 using MOSAIC (v. 1.5.0), dplyr (v. 1.1.2), and stringr (v. 1.5.0) with R (v. 4.3.0). The potential donor populations were the three European *B. taurus* populations (Holstein Friesian, Angus, and Jersey), the three African *B. taurus* populations (Muturu, Lagune, and Guinean N’Dama) and the three *B. indicus* populations (Tharparkar, Gir, and Nelore).

#### ELAI analysis

The high and low-density SNP data sets were separated and converted into BIMBAM format for each population and chromosome with PLINK (v. 1.90 beta 6.25). Local ancestry analysis was carried out for each hybrid population and autosome with 30 expectation-maximization (EM) steps, 3 upper clusters, 15 lower clusters, and 200 mixing generations using ELAI (v. 1.0) (Guan, 2014). The donor populations for each hybrid population were selected based on the results of the MOSAIC analysis.

#### Local ancestry analysis comparison

The local ancestry results were extracted using dplyr (v. 1.1.2, MOSAIC (v. 1.5.0), parallel (v. 4.3.0) (R Core Team, 2023), readr (v. 2.1.4), scales (v. 1.2.1) (Wickham *et al*, 2023c), stringr (v. 1.5.0), and tibble (v. 3.2.1) (Müller and Wickham, 2023) with R (v. 4.3.0). Mean ancestry scores across the individual hybrid animals and a genome-wide *z*-score for each of the three ancestry components were calculated for each hybrid population. Weighted mean ancestry scores and *z*-scores were calculated across the hybrid populations within each group of European hybrids, trypanotolerant African hybrids, and trypanosusceptible African hybrids for a subset of the hybrid populations selected to only include populations with a minimum of 15 animals and a relatively stable level of admixture based on a visual examination of the structure results. The local ancestry results were visualised using dplyr (v. 1.1.2), ggplot2 (v. 3.4.2), magick (v. 2.8.1) with ImageMagick (v. 6.9.12.96), magrittr (v. 2.0.3), parallel (v. 4.3.0), patchwork (v. 1.1.2), readr (v. 2.1.4), scales (v. 1.2.1), stringr (v. 1.5.0), and tibble (v. 3.2.1) with R (v. 4.3.0). Colours were generated from khroma (v. 1.10.0). The correlation between the local ancestry results from MOSAIC and ELAI were visualised using dplyr (v. 1.1.2), ggplot2 (v. 3.4.2), ggtext (v. 0.1.2), parallel (v. 4.3.0), patchwork (v. 1.1.2), readr (v. 2.1.4), and stringr (v. 1.5.0) with R (v. 4.3.0). Colours were generated from viridis (v. 0.6.3).

### Functional enrichment of introgressed regions

Functional enrichment was performed and visualised using dplyr (v. 1.1.2), ggplot2 (v. 3.4.2), ggrepel (v. 0.9.3) (Slowikowski, 2023), gprofiler2 (v. 0.2.2) (Kolberg *et al*, 2023), magick (v. 2.8.1) with ImageMagick (v. 6.9.12.96), magrittr (v. 2.0.3), parallel (v. 4.3.0), readr (v. 2.1.4), and stringr (v. 1.5.0) with R (v. 4.3.0). Colours were generated from khroma (v. 1.10.0). The background set was the set of genes within 1 Mb up-and downstream from a SNP in the high-density data set. The query sets were the genes within 1 Mb up and downstream from the SNPs with a *z*-score ≥ 2.0 for each of the ancestries.

## Results

### High-density genome-wide cattle SNP data sets

After filtering for missing genotypes (30 samples removed), identity-by-state (Figure S1; 194 samples removed), and inbreeding (Figure S2, Figure S3; 56 samples removed), there were 750 animals in the high and low-density SNP data sets (Table 1). Filtering for autosomal SNPs with a minimum call rate of 95% and MAF of at least 5% retained 614,026 SNPs in the high-density data set with a total genotyping rate of 98.20%, and 30,706 SNPs in the low-density data set with a total genotyping rate of 95.91%.

### Population genomic analyses

#### Principal component analysis

The first principal component (PC1) explained 53.59% of the of the total variation for PC1–10 in the high-density SNP data set and separated the *B. taurus* and *B. indicus* lineages (Figure 2). The second principal component (PC2) explained a further 19.71% of the total variation for PC1–10 in the high-density SNP data set and separated the European *B. taurus* and African *B. taurus* lineages (Figure 2). The hybrid animals were dispersed among the reference populations with the European hybrid animals clustering close to the European *B. taurus* group and the African hybrid animals mostly located between the African *B. taurus* and *B. indicus* groups (Figure 2). The trypanotolerant African hybrid individuals are closest to the African *B. taurus* group while the trypanosusceptible African hybrid animals are closest to the *B. indicus* group (Figure 2). The same pattern was observed for the low-density SNP data set (Figure S4).

**Figure 2.**
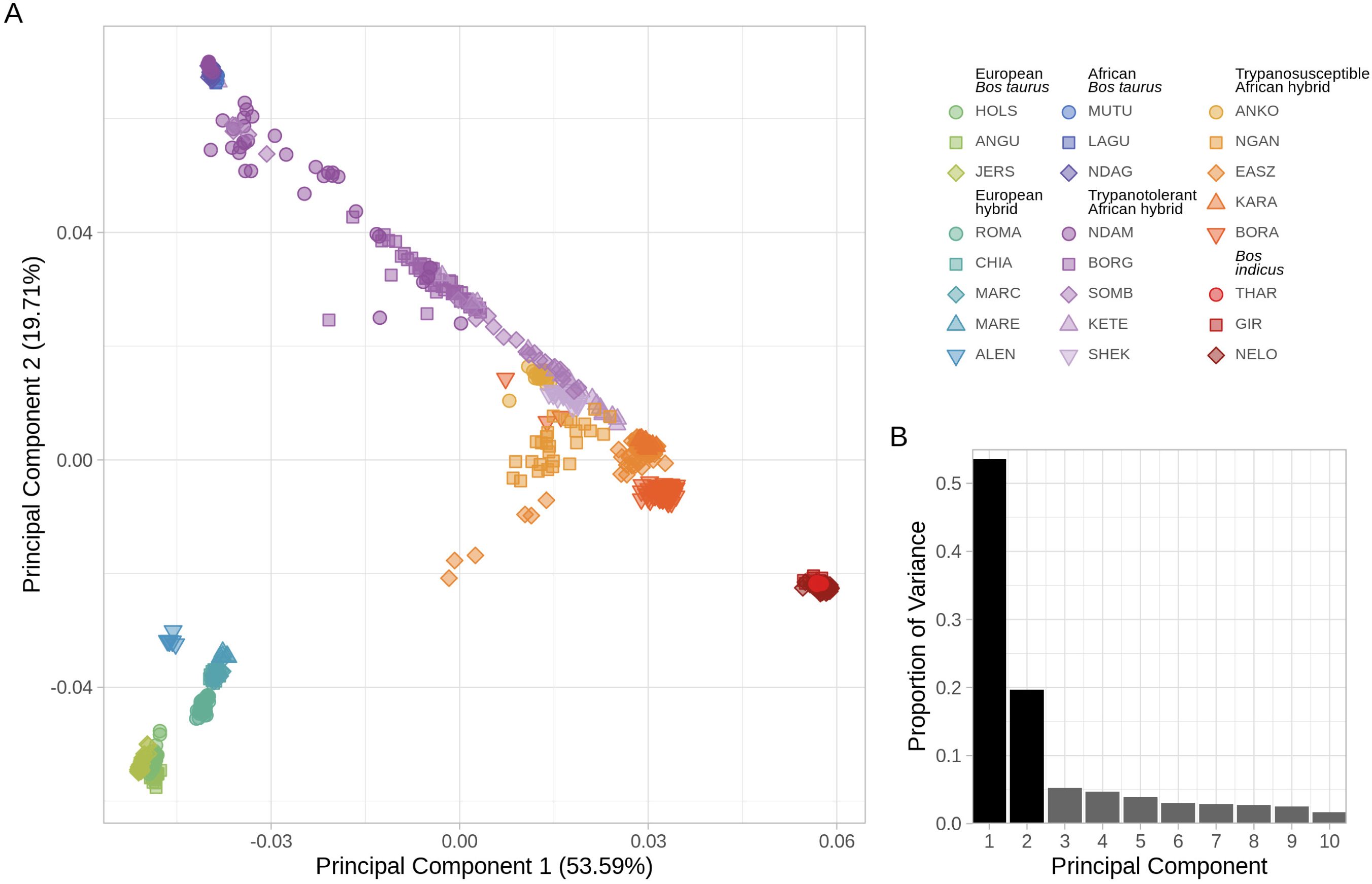
A. Principal component analysis of the selected high-density SNP data set with cattle samples coloured according to population showing the first two principal components (PC1 and PC2), and B. bar chart of the proportion of variance for the top ten PCs.

#### Genetic structure analysis

The structure results for *K* = 2 separates the *B. taurus* and *B. indicus* ancestries in the high-density SNP data set (Figure 3). With *K* = 3, the structure results divide the European and African *B. taurus* and *B. indicus* ancestries (Figure 3). For the high-density SNP data set the model complexity that maximizes marginal likelihood was 16 and the model components used to explain the structure in the data was 17 (Figure S5, Figure S6). For the low-density SNP data set the model complexity that maximizes marginal likelihood and the model components used to explain the structure in the data was 16 (Figure S7, Figure S8).

**Figure 3.**
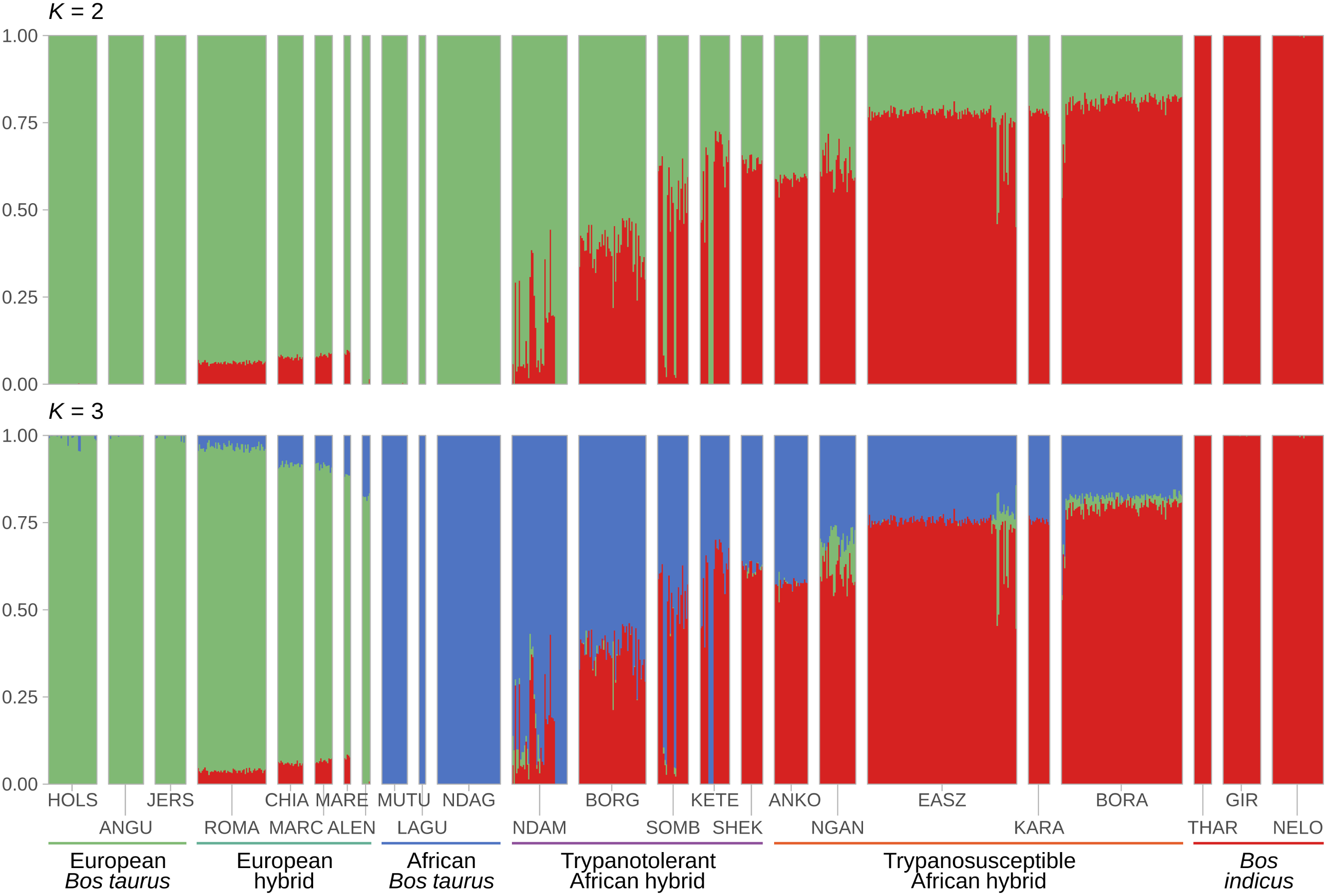
Hierarchical clustering of the high-density SNP data. Results are shown for a range of assumed values for the number of ancestral populations

#### Phylogenetic analysis

After the *B. gaurus* outgroup animals were added and filters for autosomal SNPs with a minimum call rate of 95% and MAF of at least 5% were applied there were 613,334 SNPs in the high-density SNP data set with a total genotyping rate of 99.65%, and 30,644 SNPs in the low-density SNP data set with a total genotyping rate of 99.50%. The optimum number of migration edges indicated by the first peak in *Δm* for both the high and low-density SNP data sets is three while the number of migration edges required to explain 99.8% of the variation in the data is 12 and 11 for the high and low-density SNP data sets, respectively (Figure S9, Figure S10). The phylogenetic analysis results clearly distinguish and group the European *B. taurus*, African *B. taurus*, and *B. indicus* populations into different clades with a high degree of confidence (Figure 4). The European hybrid populations are grouped with the European *B. taurus* populations with a similarly high degree of confidence, while the trypanotolerant and trypanosusceptible African hybrid populations are placed between the African *B. taurus* and *B. indicus* populations with varying degrees of confidence (Figure 4). This pattern holds regardless of the number of migration edges or SNP data set density (Figures S11–S15). The introduction of migration edges into the phylogenetic tree indicates admixture between the hybrid African and African *B. taurus* populations when *m* is set to 3 (Figure 4, Figure S14). For higher values of *m*, the admixture shown includes the European hybrid populations (Figure S12, Figure S15).

**Figure 4.**
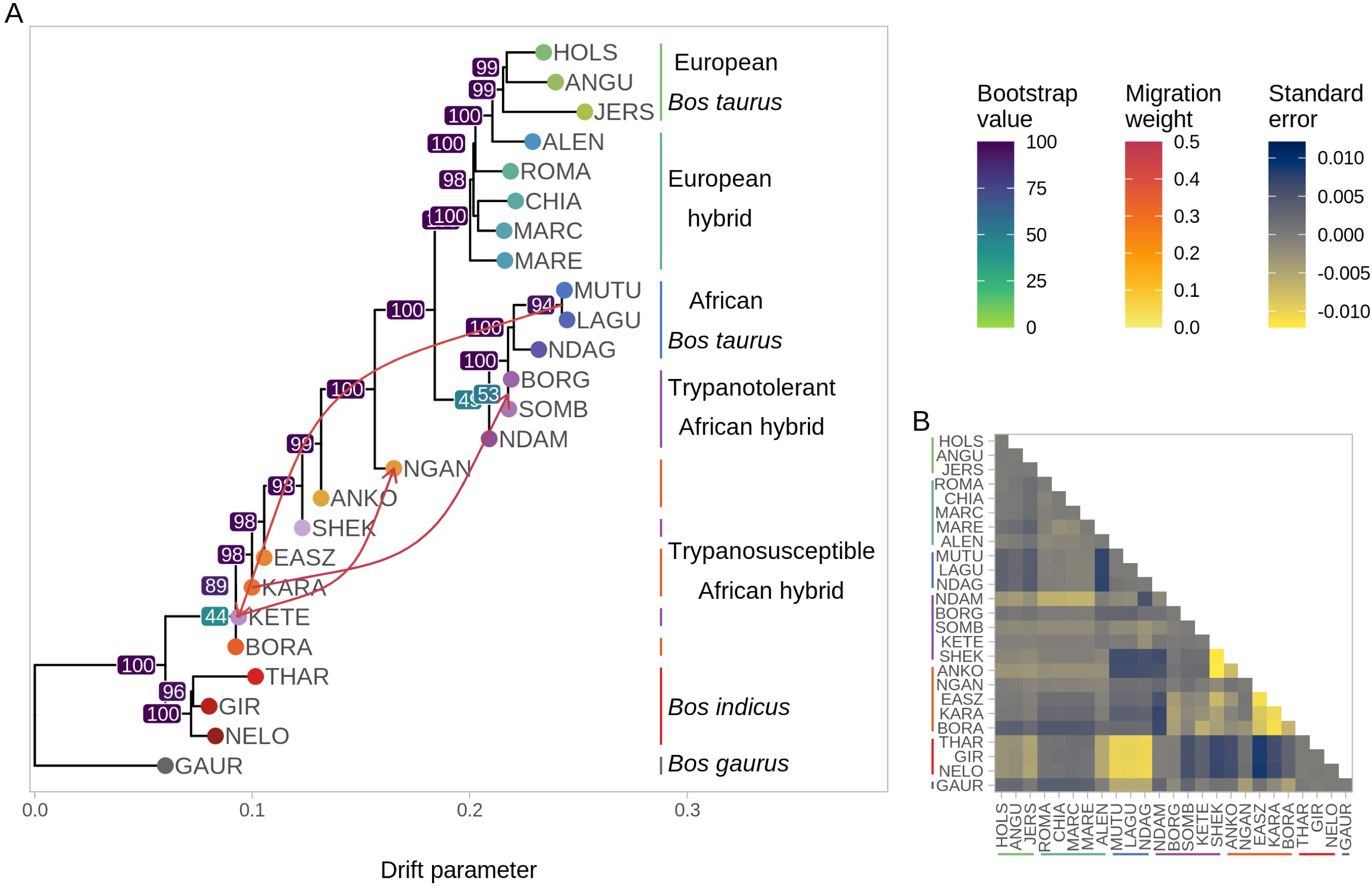
A. TreeMix phylogenetic tree for the high-density SNP data set with bootstrap values and three migration edges, and B. heatmap showing

### Local ancestry estimation

Weighted mean local ancestry results were calculated for each of the three ancestry components (European *B. taurus*, African *B. taurus*, and *B. indicus*) for each of the three hybrid groups (European hybrid, trypanotolerant African hybrid, and trypanosusceptible African hybrid) using the mean results from populations with more than 15 samples (Table 1), and relatively stable hybridisation based on visual examination of the structure results at *K* = 3 (Figure 3). The European hybrid group included the Romagnola and Chianina populations, the trypanotolerant African hybrid group included the Borgou and Sheko populations, and the trypanosusceptible African hybrid group included the Ankole, Nganda, East African Shorthorn Zebu, Karamojong, and Boran populations. The results for the high-density SNP data set showed similar patterns using both MOSAIC and ELAI when examined visually, as did the ELAI results for the low-density SNP data set (Figure 5). The MOSAIC results for the low-density SNP data set exhibited a noticeable smoothening across the genome for all three ancestry components in all three hybrid groups (Figure 5). This was particularly evident for the ancestry components with lower proportions, such as the African *B. taurus* and *B. indicus* components in the European hybrid group, and the European *B. taurus* component in the trypanotolerant African hybrid group (Figure 5). When individual chromosome results were examined, the high-density SNP local ancestry results for MOSAIC and ELAI and the low-density SNP ELAI results showed peaks for the various ancestry components around the major histocompatibility complex (MHC) located on BTA23, although this was not evident for the low-density MOSAIC results (Figure S16–S27). Correlation plots between the MOSAIC and ELAI results for each ancestry component along each chromosome indicated positive correlations for the high-density SNP results for all three hybrid groups (Figure S28–S30) while the low-density SNP results indicated much weaker or no correlations (Figure S31–S33). To identify SNPs within the peaks of local ancestry for each ancestry component genome-wide *z*-scores of the weighted mean local ancestry results were used to select SNPs with *z*-scores ≥ 2.0 for each software and data set (Table 2). There were no SNPs that passed the *z* ≥ 2 threshold for the European *B. taurus* ancestry component in the European hybrid group for the high-density SNP MOSAIC and ELAI results and the low-density SNP MOSAIC results, while the trypanotolerant and trypanosusceptible African hybrid groups had the lowest number of SNPs passing the *z* ≥ 2 threshold for the African *B. taurus* and *B. indicus* ancestry components, respectively (Table 2). Similar proportions of SNPs were found for each software and SNP data set (Table 2).

**Figure 5.**
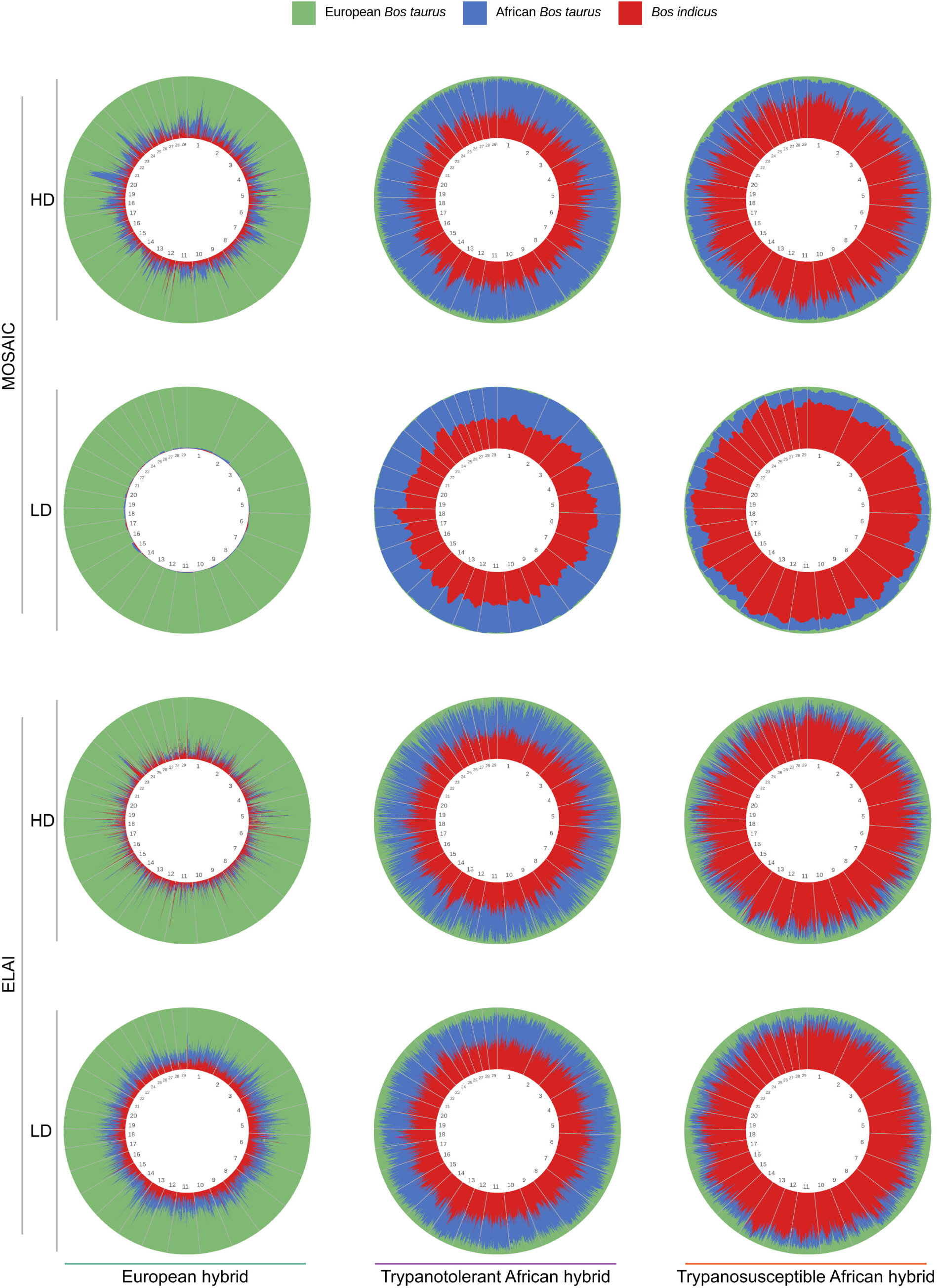
Local ancestry plots showing weighted mean European *B. taurus*, African *B. taurus*, and *B. indicus* ancestry components for European hybrid, trypanotolerant African hybrid, and trypanosusceptible African hybrid groups across all autosomes for the MOSAIC and ELAI analyses of high and low-density SNP data sets. Each vertical line on the circular genome plots represents a SNP and is coloured according to the ancestry results.

**Table 2.** Numbers of SNPs with a *z*-score ≥ 2.0 for weighted mean European *B. taurus*, African *B. taurus* and *B. indicus* ancestry components for the European hybrid, trypanotolerant African hybrid, and trypanosusceptible African hybrid groups across all autosomes for the MOSAIC and ELAI analyses of high and low-density SNP data sets. The numbers in brackets indicate the percentage of the total number of SNPs in each data set.

| Group | Ancestry | HD MOSAIC | LD MOSAIC | HD ELAI | LD ELAI |
| --- | --- | --- | --- | --- | --- |
| European hybrid | European *B. taurus* | 0<br>(0%) | 0<br>(0%) | 0<br>(0%) | 151<br>(0.49%) |
|  | African *B. taurus* | 16,511<br>(2.69%) | 1,546<br>(5.03%) | 26,561<br>(4.33%) | 1,337<br>(4.35%) |
|  | *B. indicus* | 31,674<br>(5.16%) | 1,865<br>(6.07%) | 33,110<br>(5.39%) | 1,415<br>(4.61%) |
| Trypanotolerant African hybrid | European *B. taurus* | 23,816<br>(3.88%) | 1,490<br>(4.85%) | 25,743<br>(4.19%) | 1,229<br>(4.00%) |
|  | African *B. taurus* | 7,728<br>(1.26%) | 179<br>(0.58%) | 9,056<br>(1.47%) | 627<br>(2.04%) |
|  | *B. indicus* | 18,367<br>(2.99%) | 723<br>(2.35%) | 18,083<br>(2.94%) | 801<br>(2.61%) |
| Trypanosusceptible African hybrid | European *B. taurus* | 26,035<br>(4.24%) | 1,423<br>(4.63%) | 25,083<br>(4.09%) | 1,173<br>(3.82%) |
|  | African *B. taurus* | 16,998<br>(2.77%) | 494<br>(1.61%) | 23,356<br>(3.80%) | 1,093<br>(3.56%) |
|  | *B. indicus* | 10,434<br>(1.70%) | 49<br>(1.60%) | 7,823<br>(1.27%) | 460<br>(1.50%) |
| **Total** |  | 614,026 | 30,706 | 614,026 | 30,706 |

### Functional enrichment of introgressed regions

The proportions of the numbers of genes found within 1 Mb up-and downstream from each SNP with a *z*-score ≥ 2.0 are similar to those of the numbers of SNPs found for each ancestry component in the hybrid groups for each software and SNP data set (Table 2, Table S1). There were no European *B. taurus* SNPs that passed the z ≥ 2 threshold; consequently, there were no European *B. taurus* genes for functional enrichment in the European hybrid group (Table 2, Table S1). The top driver GO terms for the African *B. taurus* genes in the European hybrid group included terms related to the MHC (*GO:0042613 MHC class II protein complex*) and other aspects of the immune system (*GO:0019882 antigen processing and presentation*, *GO:0001914 regulation of T cell mediated cytotoxicity*, and *GO:0004930 G protein-coupled receptor activity*); protein and DNA complexes and protein binding (*GO:0000786 nucleosome*, *GO:0030527 structural constituent of chromatin*, *GO:0046982 protein heterodimerization activity*, *GO:0065004 protein-DNA complex assembly*); and olfaction (*GO:0004984 olfactory receptor activity* and *GO:0050911 detection of chemical stimulus involved in sensory perception of smell*, Figure 6A). The top *B. indicus* driver GO terms also included terms relating to the MHC (*GO:0042613 MHC class II protein complex*) and other immune terms (*GO:0019882 antigen processing and presentation* and *GO:0002684 positive regulation of immune system process*), as well as cell membrane and signalling activity (*GO:0001594 trace-amine receptor activity*, *GO:0009897 external side of plasma membrane*, and *GO:0004364 glutathione transferase activity*, Figure 6A).

**Figure 6.**
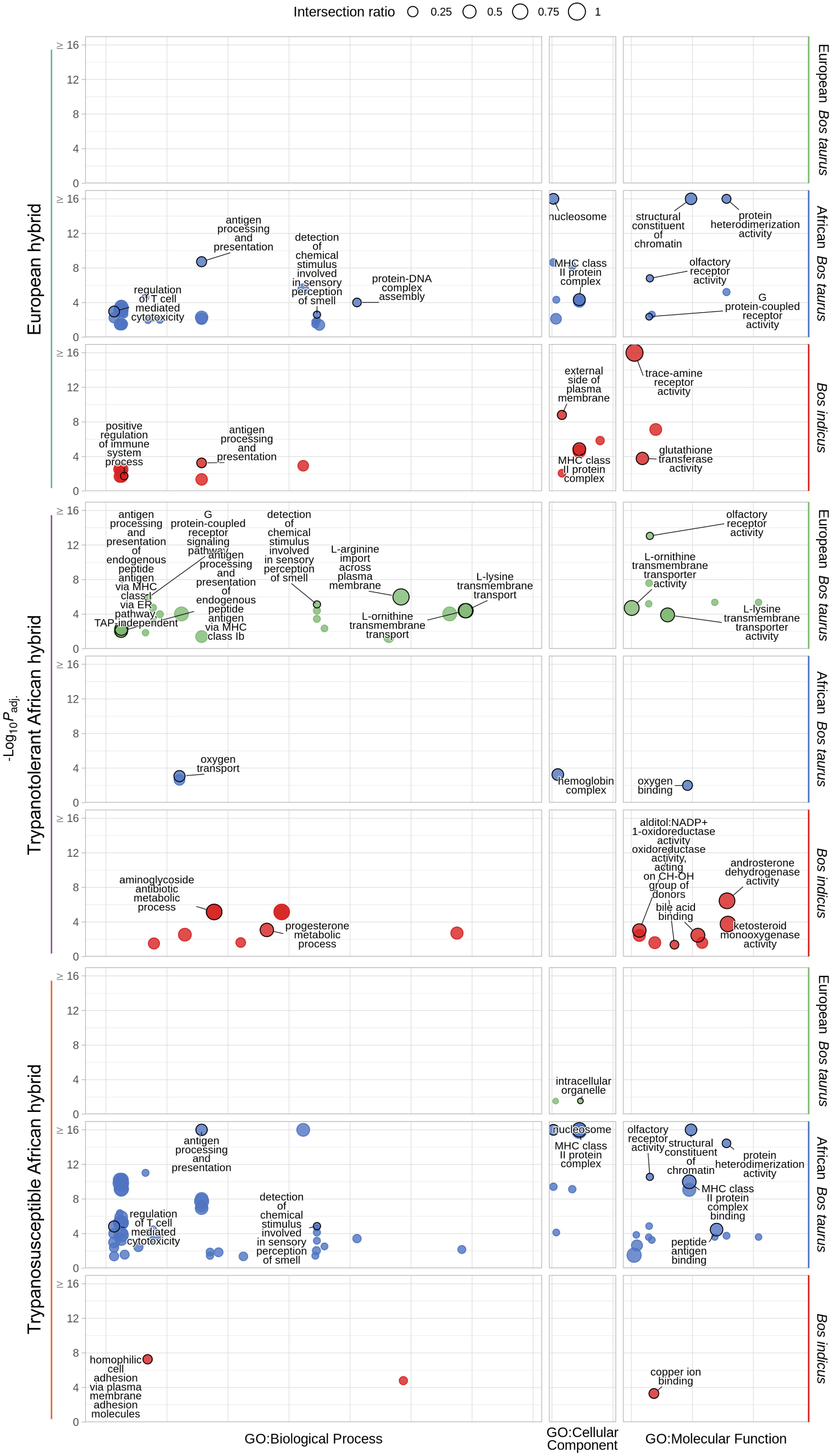
g:Profiler functional enrichment of introgressed regions in European, and trypanotolerant and trypanosusceptible African hybrid populations detected with MOSAIC and high-density SNP data. Each circle represents a significantly enriched GO term with the size indicating the ratio of the intersection between the term and the introgressed genes. The y-axis shows the −log_10_(*P*_adj_.) value and the horizontal panels and colours indicate the ancestry component. The vertical panels indicate the source of the term, and position within each panel groups terms from the same GO subtree. The top driver GO terms (up to a maximum of 10) are indicated with a black outline and label.

The trypanotolerant African hybrid group also had driver GO terms relating to the MHC (*GO:0002486 antigen processing and presentation of endogenous peptide antigen via MHC class I via ER pathway, TAP-independent* and *GO:0002476 antigen processing and presentation of endogenous peptide antigen via MHC class Ib*); other components of the immune system (*GO:0007186 G protein-coupled receptor signalling pathway*); and olfaction (*GO:0004984 olfactory receptor activity and GO:0050911 detection of chemical stimulus involved in sensory perception of smell*) among the European *B. taurus* terms (Figure 6B). In addition there were also a number of terms relating to L-amino acid transmembrane transport (*GO:0097638 L-arginine import across plasma membrane*, *GO:0000064 L-ornithine transmembrane transporter activity*, *GO:1903352 L-ornithine transmembrane transport*, *GO:1903401 L-lysine transmembrane transport*, and *GO:0015189 L-lysine transmembrane transporter activity*, Figure 6B). The top African *B. taurus* terms related to haemoglobin and oxygen binding and transport (*GO:0005833 haemoglobin complex*, *GO:0015671 oxygen transport*, *GO:0019825 oxygen binding*), while the top *B. indicus* terms related to metabolic processes (*GO:0047023 androsterone dehydrogenase activity*, *GO:0030647 aminoglycoside antibiotic metabolic process*, *GO:0047086 ketosteroid monooxygenase activity*, *GO:0042448 progesterone metabolic process*, *GO:0004032 alditol:NADP+ 1-oxidoreductase activity*, *GO:0032052 bile acid binding*, and *GO:0016614 oxidoreductase activity, acting on CH-OH group of donors*, Figure 6B).

For the trypanosusceptible African hybrid group the only driver GO term for the European *B. taurus* ancestry component related to intracellular organelles (*GO:0043229 intracellular organelle*, Figure 6C). The top African *B. taurus* terms included those related to the MHC (*GO:0042613 MHC class II protein complex*, and *GO:0023026 MHC class II protein complex binding*); other components of the immune system (*GO:0019882 antigen processing and presentation*, *GO:0001914 regulation of T cell mediated cytotoxicity*, and *GO:0042605 peptide antigen binding*); olfaction (*GO:0004984 olfactory receptor activity*, and *GO:0050911 detection of chemical stimulus involved in sensory perception of smell*); and protein-DNA complex and protein binding (*GO:0030527 structural constituent of chromatin*, *GO:0000786 nucleosome*, *GO:0046982 protein heterodimerization activity*, Figure 6C). The top *B. indicus* terms included cell adhesion (*GO:0007156 homophilic cell adhesion via plasma membrane adhesion molecules*) and metal ion binding (*GO:0005507 copper ion binding*, Figure 6C).

Similar GO term enrichment patterns were also observed using low-density SNP data with MOSAIC (Figure S34), and for ELAI with both high and low-density SNP data (Figure S35, Figure S36).

## Discussion

The results of the population genomic analyses in hybrid European and African cattle populations are consistent with previously published studies that have used modest numbers of genetic markers (e.g., microsatellites) and genome-wide SNP data (Barbato *et al*, 2020; Decker *et al*, 2014; Hanotte *et al*, 2002; Kim *et al*, 2020; MacHugh *et al*, 1997; Ward *et al*, 2022). Visualisation of PCA results by plotting PC1 and PC2 recovered the classic “*Bos* triangle” with the first two PCs explaining a very high proportion of the total variation for PC1–10 within the data (73.30%). PC1 and PC2 separated the reference European *B. taurus*, African *B. taurus*, and *B. indicus* populations with the hybrid animal samples dispersed within the triangle with locations determined by three-way global admixture proportions (Figure 2). The locations of the various hybrid populations nearer to the reference populations they share the most ancestry with is in agreement with previous studies (Bahbahani *et al*, 2017; Barbato *et al*, 2020; Upadhyay *et al*, 2017; Verdugo *et al*, 2019; Ward *et al*, 2022; Wragg *et al*, 2022). In addition, the clustering of some of the animals in the African trypanotolerant hybrid populations with the African *B. taurus* reference populations indicate that some of these animals have very high levels of African taurine ancestry (Figure 2). In this regard, it is important to note that although a diverse panel of European *B. taurus*, African *B. taurus*, *B. indicus*, and hybrid cattle in the design and validation of the BovineHD 777K BeadChip (Illumina, 2015), ascertainment bias may affect the placement of hybrid cattle in a PCA plot (Dokan *et al*, 2021; McTavish and Hillis, 2015). However, genome-wide multi-locus dimension reduction tools are typically substantially less affected by ascertainment bias than analyses such as estimation of diversity statistics such as the fixation index (*F*ST) or selection signal detection, which use individual SNP locus frequency-based statistics (Albrechtsen *et al*, 2010; Malomane *et al*, 2018; Porto Neto and Barendse, 2010).

The results of the genetic structure analysis for *K* = 2 and *K* = 3 mirror those of the PCA with the first major split evident for the *B. taurus* and *B. indicus* populations and the second split separating the African and European *B. taurus* populations (Figure 3). The locations of animals in the hybrid populations reflect global admixture proportions that are in agreement with both their positions on the PCA and previous studies (Bahbahani *et al*, 2017; Barbato *et al*, 2020; Upadhyay *et al*, 2017; Verdugo *et al*, 2019; Ward *et al*, 2022; Wragg *et al*, 2022) (Figure 2). The number of modelled *K* values that best explain the variation among the 24 populations examined in the study was 16–17, indicating that some of the populations are closely related to the point that they may not be genetically distinct discrete populations (Raj *et al*, 2014). The genetic structure results also show the variation within the hybrid populations in terms of global admixture (Figure 3). Some populations, such as the European and trypanosusceptible African hybrid groups, show a relatively consistent level of global admixture across each population while the trypanotolerant African hybrid group is more variable (Figure 3). This indicates that the European and trypanosusceptible African hybrid breeds are more long-established hybrid populations (“stable crossbreds”) and that the hybridisation within the African trypanotolerant hybrid populations is more recent and dynamic. Some of the more extreme examples, such as the N’Dama hybrid, Somba, and Keteku populations, indicate that some animals are not hybrids and are instead pure African *B. taurus* (Figure 3). This is also in agreement with the PCA results and is likely due to the origins of the samples from a range of studies that sampled animals from different populations that were classified as the same breed or breed subtype (Bahbahani *et al*, 2017; Barbato *et al*, 2020; Upadhyay *et al*, 2017; Verdugo *et al*, 2019; Ward *et al*, 2022; Wragg *et al*, 2022) (Figure 2).

The results of the phylogenetic analysis are also in agreement with the PCA and genetic structure results (Figure 4). The reference populations are unambiguously separated into the expected groupings (*European B. taurus*, African *B. taurus*, and *B. indicus*) with bootstrap values of 99–100, as are the European hybrid populations (Figure 4). The trypanotolerant and trypanosusceptible African hybrid populations are spread between the African *B. taurus* and *B. indicus* reference populations, with some hybrid branch clusters exhibiting low bootstrap values, indicating instability in the clade structure because of taurine/indicine admixture (Figure 4). This is also where the strongest gene flow events are inferred as modelled migration edges, demonstrating the higher levels of indicine admixture in the trypanotolerant and trypanosusceptible African hybrid populations compared to the European hybrid populations (Figure 4).

The local ancestry results show similar patterns of peaks and troughs dispersed across the genome for each LAI software tool used (MOSAIC and ELAI), the hybrid cattle group examined (European hybrid, trypanotolerant African hybrid, and trypanosusceptible African hybrid), and each genome-wide SNP data set analysed (low or high-density); the exception being the low-density SNP data set results obtained using MOSAIC (Figure 5). This may be due to the rephasing algorithm implemented by default as part of the MOSAIC analysis, which may overcorrect for phasing errors in low-density SNP data set (Salter-Townshend and Myers, 2019). Alternatively, the smoothened results may be due to difficulty automatically estimating the age of admixture with low-density SNP data in MOSAIC (Salter-Townshend and Myers, 2019). Comparing the MOSAIC high-density SNP data set results with the ELAI results using both the high and low-density SNP data sets (Figure 5), the similar genome-wide patterns of local ancestry obtained indicate that, despite the differences in the software used, there are robust signals of local ancestry discernible in these populations. This is evident in the marked three-way ancestry diversity around the MHC region on BTA23 seen in these results (Figures S16–S27), particularly for the European hybrid group using both MOSAIC and ELAI and the high-density SNP data set where there is a clear signature of elevated African taurine and indicine ancestry (Figure S16, Figure S22). This tendency for increased genomic introgression in the bovine MHC region is likely a consequence of the balancing selection that maintains high MHC gene polymorphism due to the key function of MHC class I and II proteins in presentation of antigenic peptides from rapidly evolving pathogens to CD4^+^ and CD8^+^ T cells and via interactions with receptors on natural killer (NK) cells (Codner *et al*, 2012; Ellis, 2004; Ellis and Hammond, 2014). Balancing selection acting on pre-existing trans-specific polymorphisms and introgressed variants would give rise to extensive polymorphism in the MHC region (Radwan *et al*, 2020) and this has been observed in several species (Hedrick, 2013), including *Homo sapiens* where there is evidence that Neanderthal (*Homo sapiens neanderthalensis*) and Denisovan (*Homo sapiens* subsp. ‘Denisova’) MHC gene variants have readily introgressed into anatomically modern human populations (Liston *et al*, 2021; Racimo *et al*, 2015). However, it is also important to note that there are known difficulties in genotyping the MHC region (Dicks *et al*, 2021).

The correlations observed between the MOSAIC and ELAI results for the high-density SNP data sets (Figures S28–S33) provides additional evidence that supports the visually apparent similarities in the local ancestry signals observed across the bovine genome using both software approaches. The proportions of the numbers of SNPs passing the genome-wide threshold (*z*-score ≥ 2.0) were similar for the MOSAIC and ELAI analyses using both the high and low-density SNP data sets, which despite the visual differences in the local ancestry results, indicates that similar numbers of ancestry peaks can be detected using the *z*-score approach (Table 2). The lack of SNPs passing the threshold for the European *B. taurus* ancestry component in the European hybrid group is likely due to the high and relatively uniform proportion of the European *B. taurus* ancestry component across the genome. This would give rise to a situation such that no SNPs could pass the threshold of two standard deviations from the mean (Figure 5). Similarly, the lower numbers of SNPs passing the threshold for the African *B. taurus* and *B. indicus* ancestry components for the trypanotolerant and trypanosusceptible African hybrid groups, respectively, is likely due to the higher proportions of the reference population ancestries to which each hybrid group is most closely related (Figure 5).

The introgressed genomic regions for the three hybrid population samples show several distinct patterns in terms of functional enrichment. All three hybrid groups had significant driver GO terms relating to the MHC (Figure 6), which directly encompass MHC genes (e.g., MHC class I and II) and other genes encoding proteins that interact with MHC gene products. Visual examination of the local ancestry results supports this observation as do previous LAI studies in cattle (Figure 5, Figures S16– S27) (Buggiotti *et al*, 2021; Chen *et al*, 2020; Guan *et al*, 2022; Li *et al*, 2023). Other immune system related driver GO terms were also found to be significant for the three hybrid groups. Several of these terms contain genes that are either up or downstream from MHC genes in biological pathways, underscoring the importance of MHC-related immunobiology in admixed cattle. More generally, it is notable that immune genes are well represented in the top functional enrichment categories for the introgressed genomic regions since there are well documented differences among European *B. taurus*, African *B. taurus*, and *B. indicus* cattle populations in terms of susceptibilities to various infectious diseases such as bovine tuberculosis caused by *Mycobacterium bovis* (Allen *et al*, 2010; Lee *et al*, 2024); East Coast fever and tropical theileriosis caused by *Theileria parva* and *Theileria annulate*, respectively (Bahbahani and Hanotte, 2015); and AAT caused by *Trypanosoma* spp. (Yaro *et al*, 2016). In this regard, many of the genes highlighted by LAI through retention of taurine ancestry in the trypanotolerant African hybrid population may represent putative candidate genes underlying the multigenic trypanotolerance trait. For example, in this group, genes associated with haemoglobin, and oxygen binding and transport cellular processes were highlighted by the GO term functional enrichment for retained African *B. taurus* genomic ancestry (Figure 6). This may reflect positive selection of genomic variants that enhance control of anaemia, which is understood to be a key feature of the trypanotolerance trait in cattle (Kambal *et al*, 2023).

Driver GO terms relating to olfaction were also significantly enriched across the three hybrid cattle groups (Figure 6). Genes related to olfaction, such as olfactory receptor (OR) genes, have been identified in previous functional population genomics analyses of admixed cattle populations with taurine and indicine ancestry. These include, for example, genes containing breed-specific missense SNPs in admixed Ethiopian cattle (Zegeye *et al*, 2023), genes within genomic regions with evidence for selection signatures in admixed Turkish and Chinese cattle (Demir *et al*, 2023; Sun *et al*, 2023), and genes in population-differentiated copy-number variation regions (CNVRs) in African hybrid cattle (Jang *et al*, 2021). This may be due to the relatively large number of OR genes dispersed across the cattle genome, which, at more than 800 functional loci is comparable to the OR gene repertoire in the domestic dog (*Canis familiaris*) (Lee *et al*, 2013; Niimura and Nei, 2007). However, recent studies have suggested that more than 500 olfactory receptors may be expressed by macrophages, immune cells involved in detection and phagocytosis of pathogens (Orecchioni *et al*, 2022). Macrophages are the host’s first line of defence to mycobacterial infections with evasion and reprogramming of host macrophages being key components of host-pathogen interaction (Hall *et al*, 2024). In this regard, it is therefore noteworthy that sequence variation at olfactory receptor gene loci has been shown to be associated with susceptibility to *M. bovis* infection in cattle (Ring *et al*, 2019).

An alternative hypothesis for enrichment of olfaction-related genes, however, could relate to detection of odorants associated with MHC diversity and selection of mates (Santos *et al*, 2010; Ziegler *et al*, 2010), although this is unlikely to be a major factor in managed male-biased cattle husbandry systems. Similarly, cattle populations under intensive human control and management are unlikely to require a keen sense of smell to find food and avoid danger; however, introgressive natural selection is likely to be acting on olfaction-related genes in free-ranging admixed African cattle populations exposed to a wide range of environmental and predation challenges (Mwai *et al*, 2015).

The comparative LAI analyses we have performed using low and high-density SNP array data sets in various groups of admixed cattle with taurine and indicine genomic ancestry provides a framework for applying LAI to much larger data sets that will encompass millions of SNPs. In addition, our results will provide a context for understanding the genomic basis of heterosis in admixed cattle, particularly as it dissipates beyond the F_1_ generation (Syrstad, 1985). Also, identification of genomic regions that have been subject to introgressive selection will provide important information for genome-enabled breeding in admixed cattle populations, particularly in Africa (Marshall *et al*, 2019; Mrode *et al*, 2019). Finally, the methodologies that we describe here can be applied to other hybrid cattle populations, for example, admixed breeds in Anatolia and the Middle East that have had much longer histories of taurine/indicine genetic exchange (Verdugo *et al*, 2019).

## Acknowledgements

We thank Morris Agaba, Olivier Hanotte, Stephen J. Kemp, John A. Browne, Daniel G. Bradley, and Stephen V. Gordon for assistance with sample resources and for useful scientific discussion. This research work was funded by Science Foundation Ireland (SFI) under Investigator Programme Awards (grant nos: SFI/01/F.1/B028 and SFI/15/IA/3154). JAW was supported by the Centre for Research Training in Genomics Data Science (grant no. SFI/18/CRT/6214).

## Author contribution statement

GPM was responsible for analysis, data curation, lab work, interpretation of results, study design, visualisation, and writing - original draft. JAW was responsible for data provision, lab work, interpretation of results, and writing - review & editing. SIN was responsible for data provision, interpretation of results, and writing - review & editing. LAF was responsible for data provision, interpretation of results, and writing - review & editing. MST was responsible for interpretation of results, software provision, and writing - review & editing. EWH was responsible for lab work, sample collection and provision, and writing - review & editing. GMO was responsible for lab work, sample collection and provision, and writing - review & editing. KGM was responsible for lab work, sample collection and provision, and writing - review & editing. TJH was responsible for guidance and writing - review & editing. DEM was responsible for data provision, funding acquisition, lab work, interpretation of results, sample collection and provision, study design, supervision, and writing - original draft.

## Conflict of Interest

The authors declare no competing interests.

## Data archiving

New Illumina^®^ BovineHD 777K BeadChip SNP data sets generated for this study have been deposited in the Dryad data repository at doi.org/10.5061/dryad.w3r22810n. The computer code required to repeat and reproduce the analyses is available at doi.org/10.5281/zenodo.11491949.

## Research Ethics Statement

For this study new Illumina^®^ BovineHD 777K BeadChip SNP data sets were generated for 39 individuals (23 Somba, 8 N’Dama and 8 Boran). The Somba individuals were obtained from DNA samples that were previously published as part of microsatellite-based surveys of cattle genetic diversity in the early 1990s and the N’Dama and Boran individuals were obtained from unpublished DNA samples collected during a time-course infection experiment carried out in 2003. This livestock DNA sampling work was completed prior to the requirement for Institutional Permission in Ireland, which is based on European Union Directive 2010/63/EU; however, all efforts were made to ensure ethical handling of all animal subjects.

## Notes

https://doi.org/10.5061/dryad.w3r22810n

https://doi.org/10.5281/zenodo.11491949

### Competing Interest Statement

The authors have declared no competing interest.

