## Supplementary Material for "Genome-wide local ancestry and the functional consequences of admixture in African and European cattle populations"

**Table S1.** Numbers of genes within 1 Mb up- and downstream of SNPs with a *z*-score ≥ 2.0 for weighted mean European *Bos taurus*, African *Bos taurus* and *Bos indicus* ancestry components for the European hybrid, trypanotolerant African hybrid, and trypanosusceptible African hybrid groups across all autosomes for the MOSAIC and ELAI analyses of high- and low-density SNP data sets. The numbers in brackets indicate the percentage of the total number of SNPs in the data set.

| **Group** | **Ancestry** | **HD MOSAIC** | **LD MOSAIC** | **HD ELAI** | **LD ELAI** |
|---|---|---|---|---|---|
| European hybrid | European *B. taurus* | 0 (0%) | 0 (0%) | 0 (0%) | 20 (0.08%) |
| | African *B. taurus* | 720 (2.76%) | 1,980 (7.61%) | 585 (2.24%) | 457 (1.76%) |
| | *B. indicus* | 1,131 (4.33%) | 1,564 (6.01%) | 954 (3.66%) | 719 (2.76%) |
| Trypanotolerant African hybrid | European *B. taurus* | 538 (2.06%) | 689 (2.65%) | 382 (1.46%) | 285 (1.10%) |
| | African *B. taurus* | 271 (1.04%) | 191 (0.73%) | 11 (0.04%) | 192 (0.74%) |
| | *B. indicus* | 976 (3.74%) | 1,117 (4.29%) | 556 (2.13%) | 333 (1.28%) |
| Trypanosusceptible African hybrid | European *B. taurus* | 540 (2.07%) | 584 (2.24%) | 250 (0.96%) | 372 (1.43%) |
| | African *B. taurus* | 615 (2.36%) | 168 (0.65%) | 268 (1.03%) | 352 (1.35%) |
| | *B. indicus* | 451 (1.73%) | 686 (2.64%) | 216 (0.83%) | 214 (0.82%) |
| | **Total** | 26,101 | 26,017 | 26,101 | 26,017 |

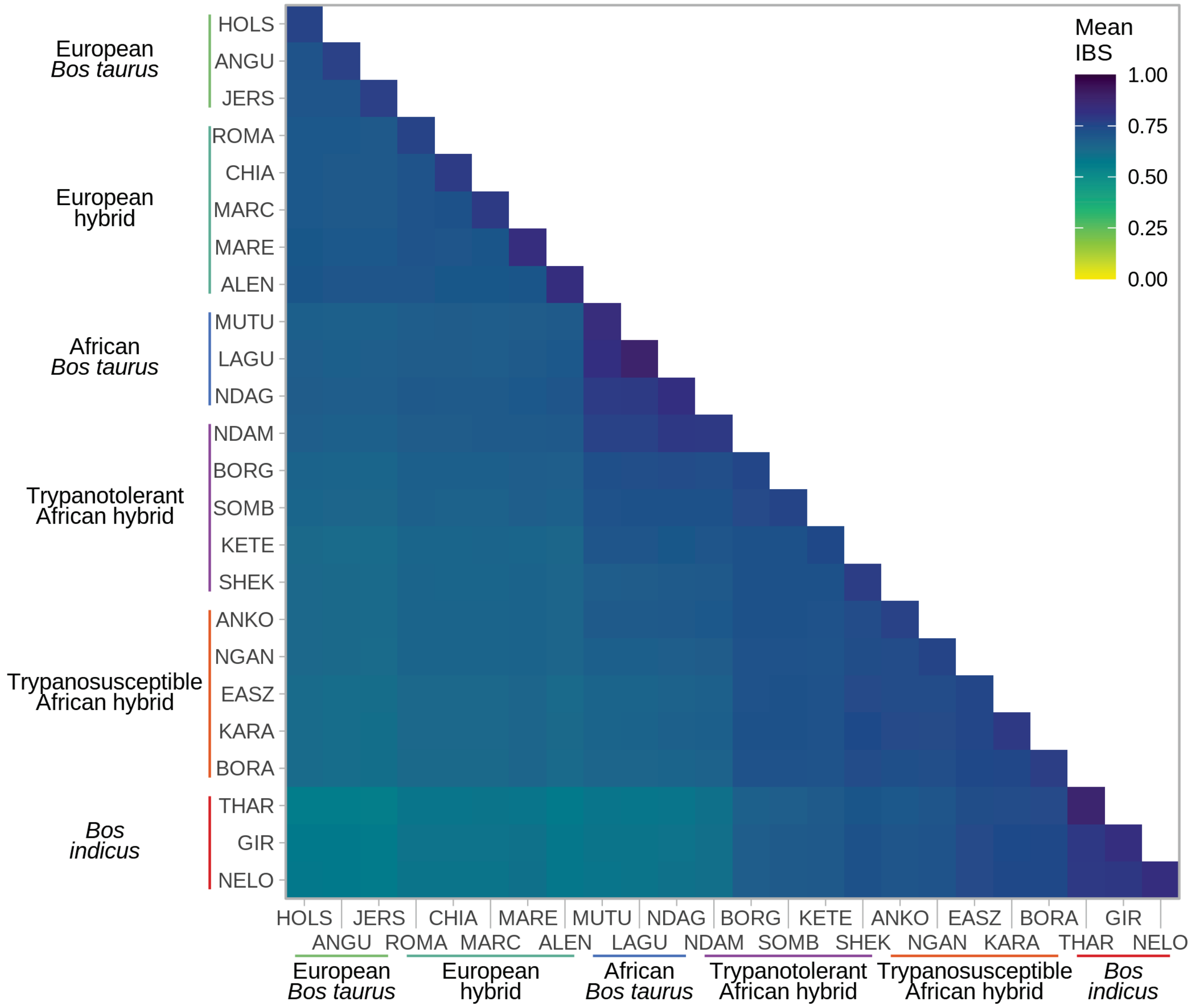


**Figure S1.** Heatmap of mean identity-by-state values for the high-density SNP data set.

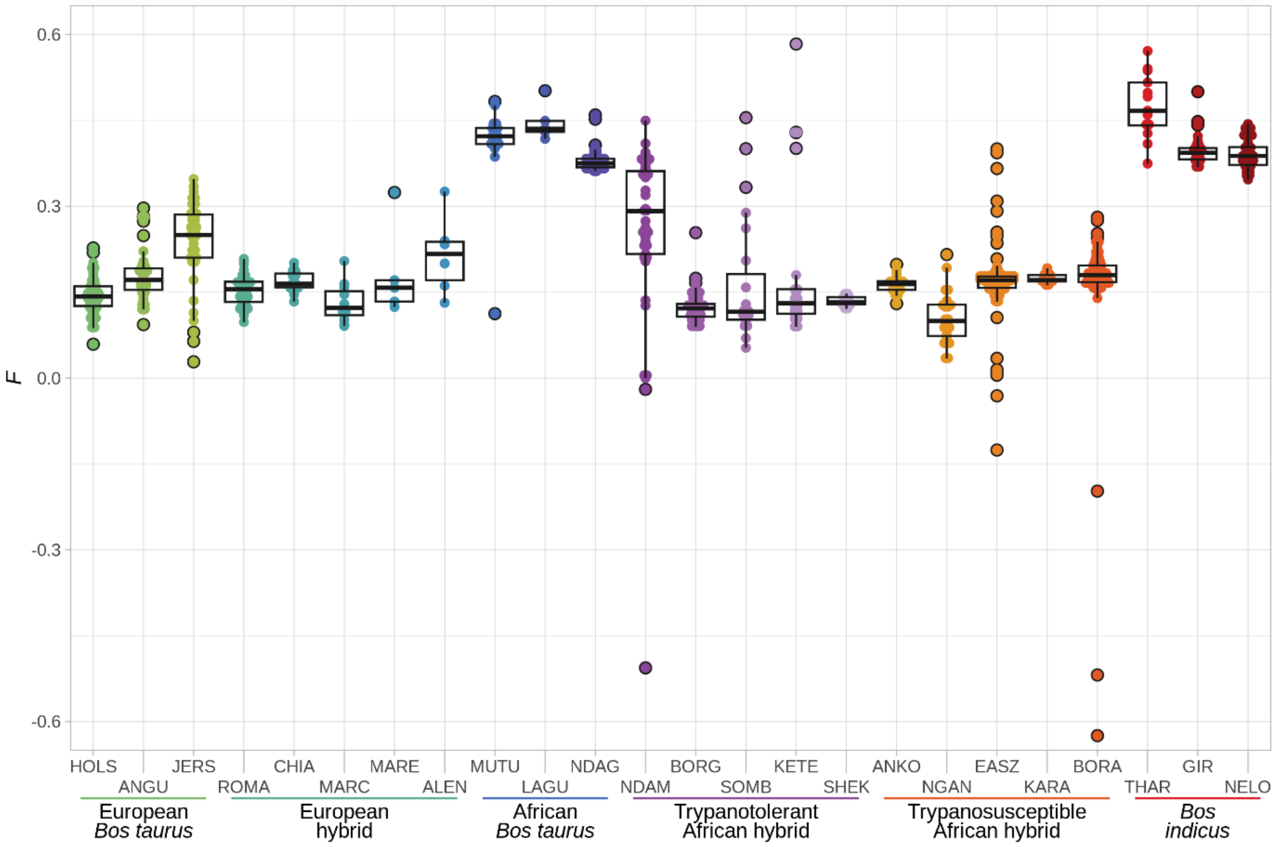


**Figure S2.** Tukey box plots showing the distribution of inbreeding values (*F*) for the high-density SNP data for each population. Outliers are indicated with a black outline.

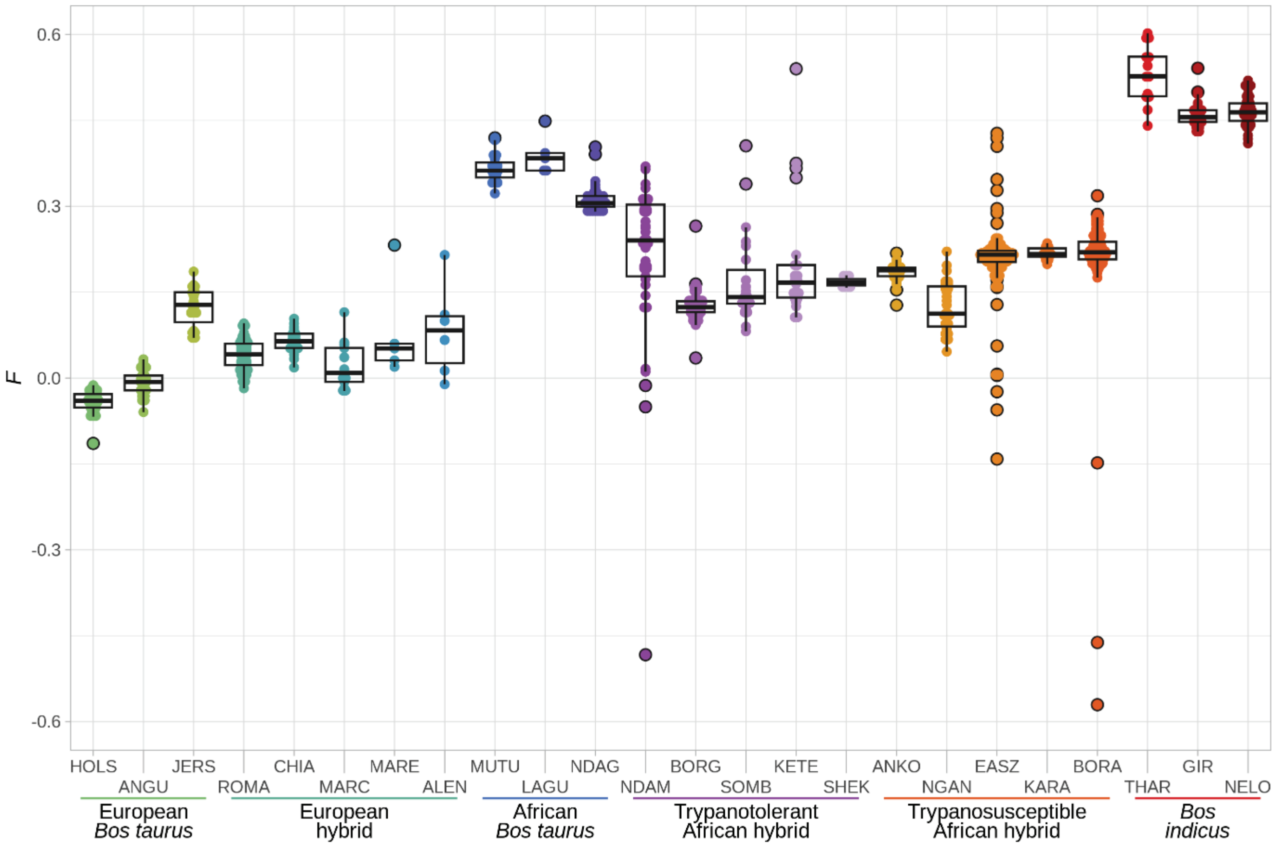


**Figure S3.** Tukey box plots showing the distribution of inbreeding values (*F*) for the low-density SNP data for each population after inbreeding filters were applied. Outliers are indicated with a black outline.

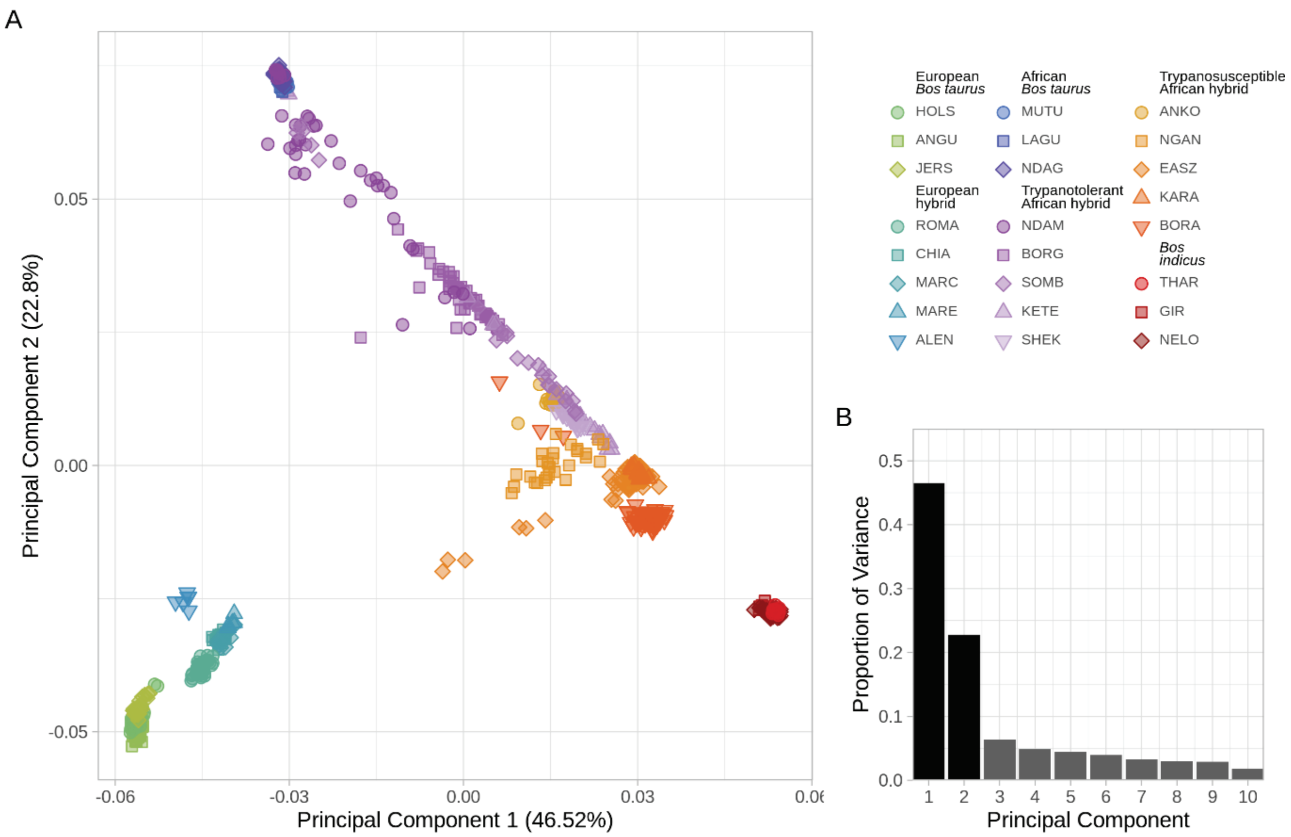


**Figure S4.** A. Principal component analysis of the selected low-density SNP data set with cattle samples coloured according to population showing the first two principal components (PC1 and PC2), and B. bar chart of the proportion of variance for the top ten PCs.

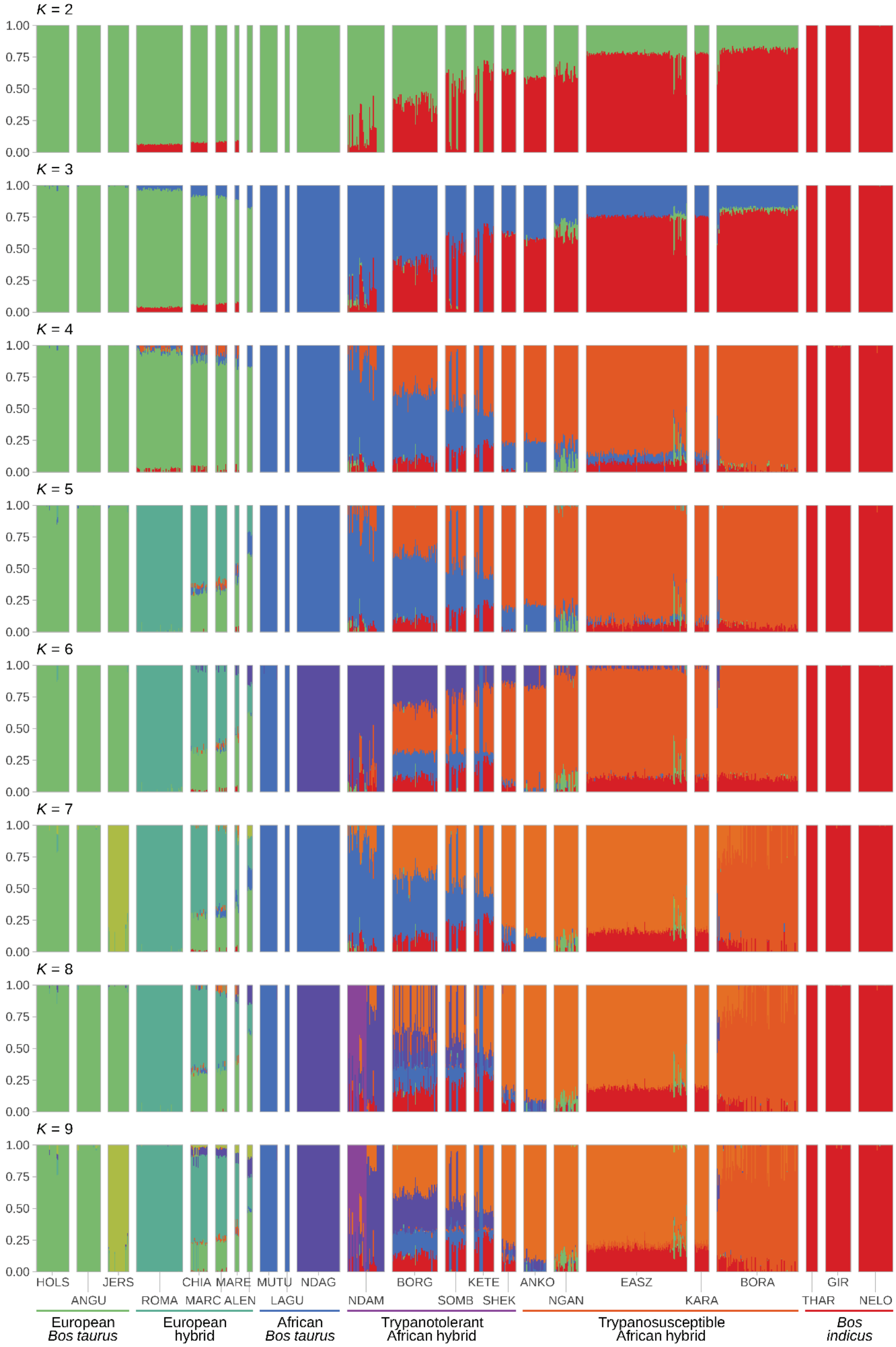


**Figure S5.** Hierarchical clustering of cattle samples using the high-density SNP data set. Results are shown for a range of assumed values for the number of ancestral populations ($K$ = 2–9).

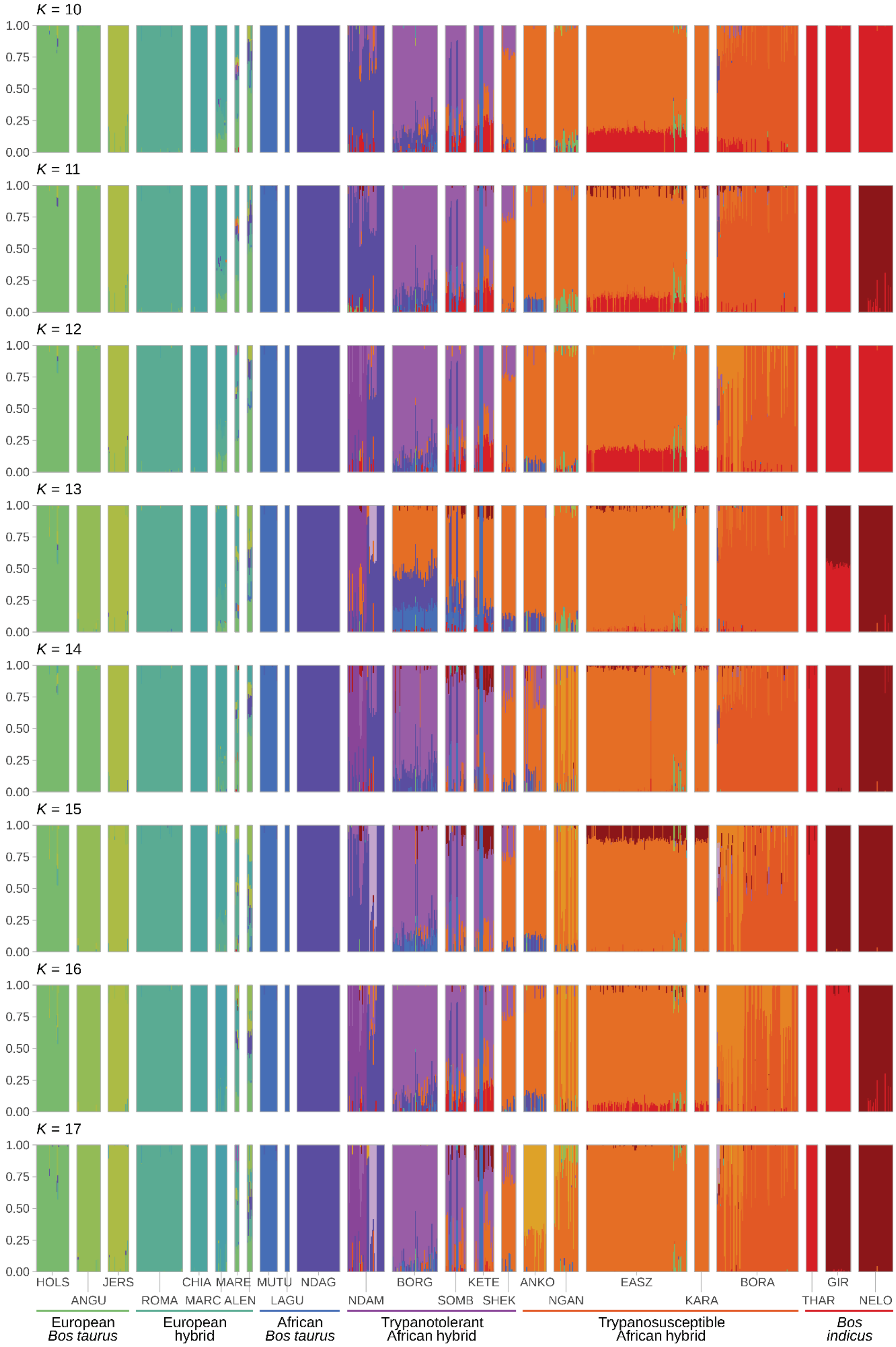


**Figure S6.** Hierarchical clustering of cattle samples using the high-density SNP data set. Results are shown for a range of assumed values for the number of ancestral populations ($K$ = 10–17).

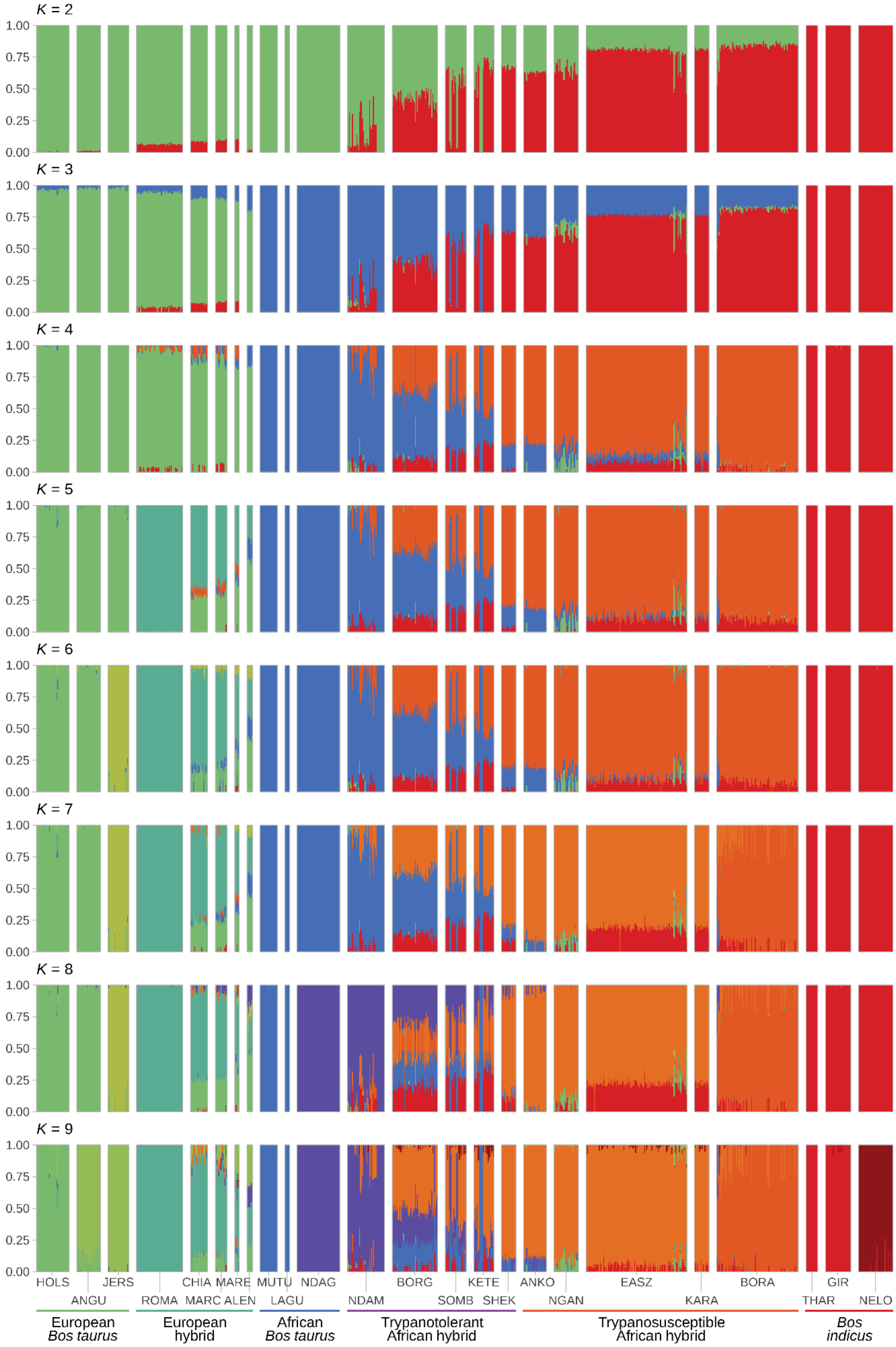


**Figure S7.** Hierarchical clustering of cattle samples using the low-density SNP data set. Results are shown for a range of assumed values for the number of ancestral populations ($K$ = 2–9).

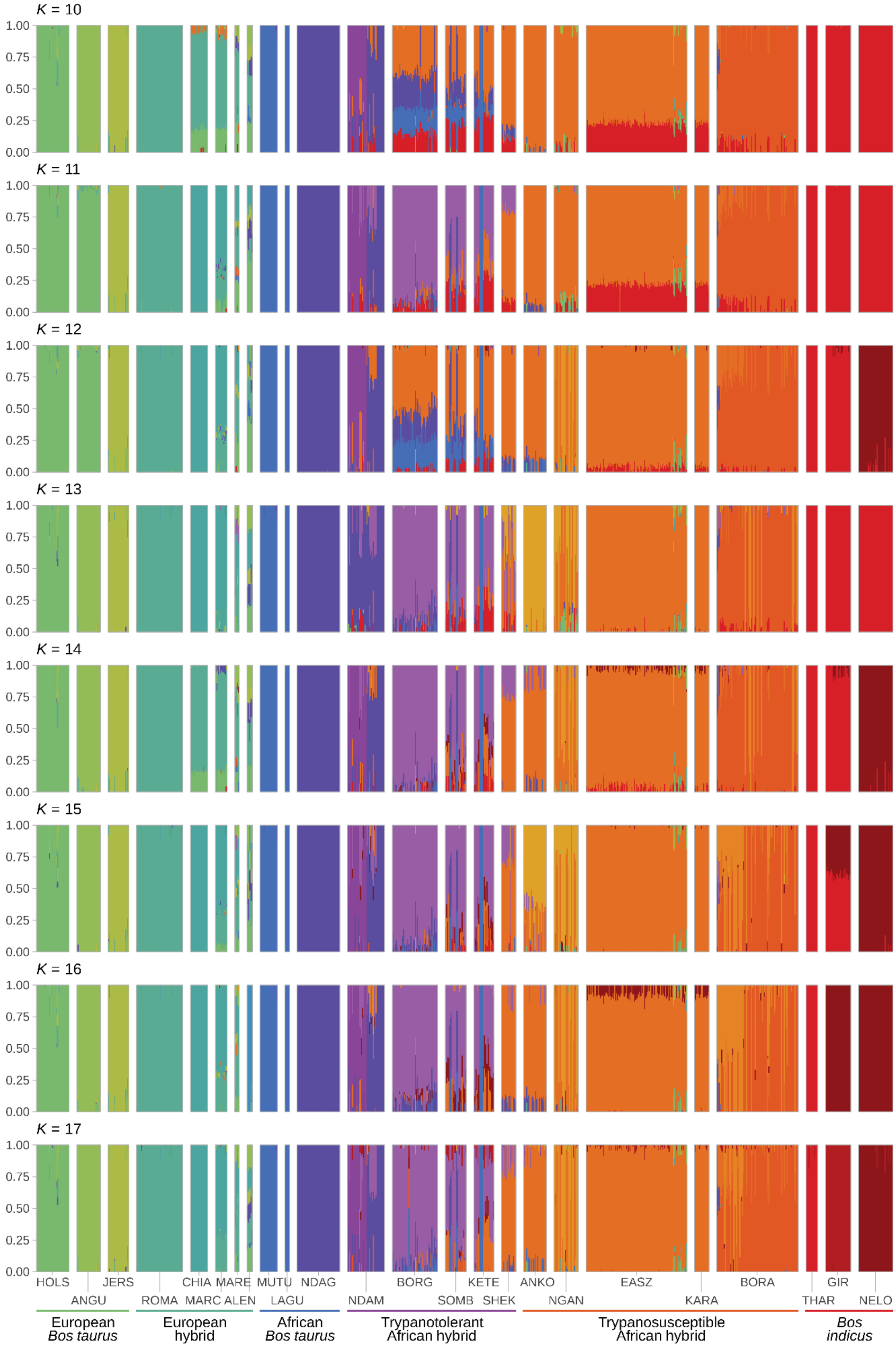


**Figure S8.** Hierarchical clustering of cattle samples using the low-density SNP data set. Results are shown for a range of assumed values for the number of ancestral populations ($K$ = 10–17).

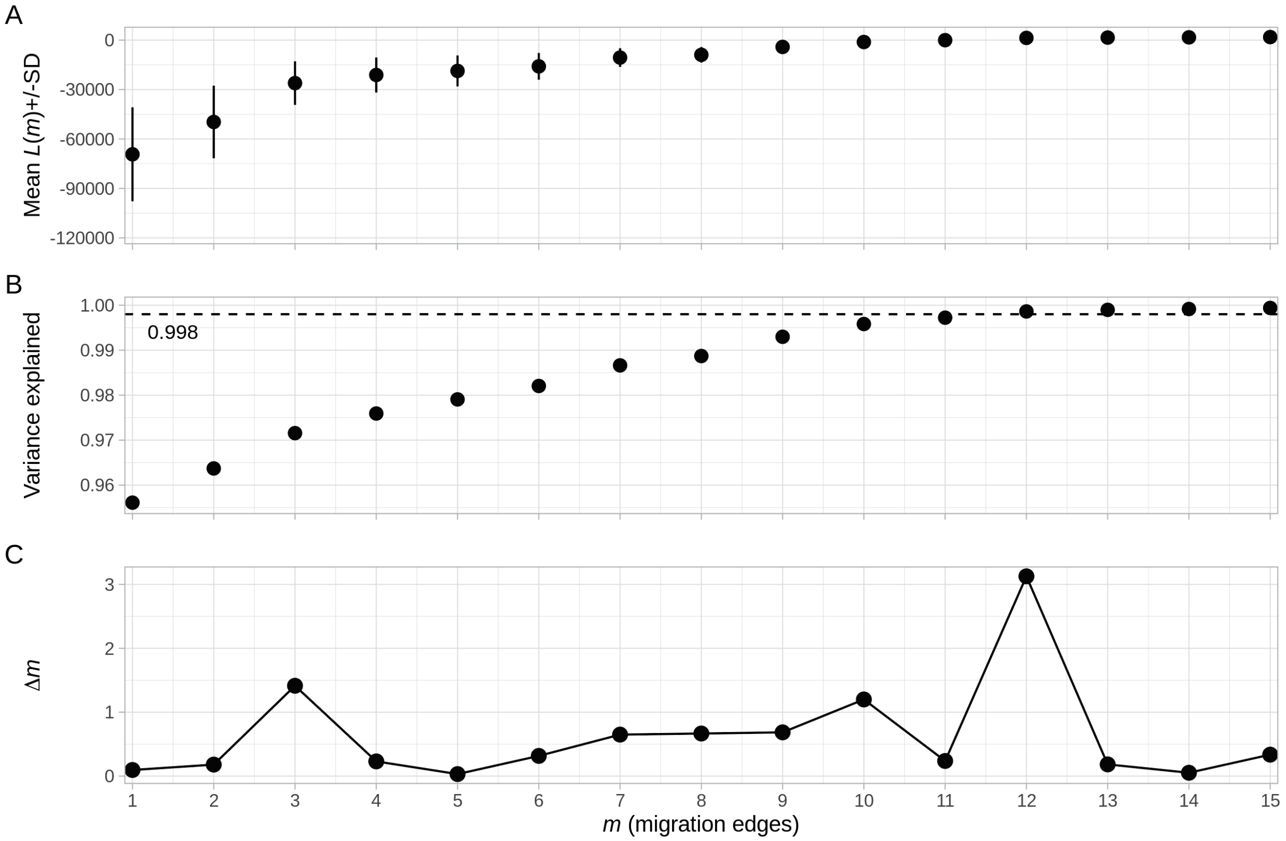


**Figure S9.** OptM results for the high-density SNP data set. **A.** the mean and standard deviation (SD) across 10 iterations for the composite likelihood (*L*(*m*)). **B.** the proportion of variance explained showing the 99.8% threshold (horizontal dotted line) recommended by Pickrell and Pritchard (2012). **C.** the second-order rate of change (*Δm*) across migration edges (*m*).

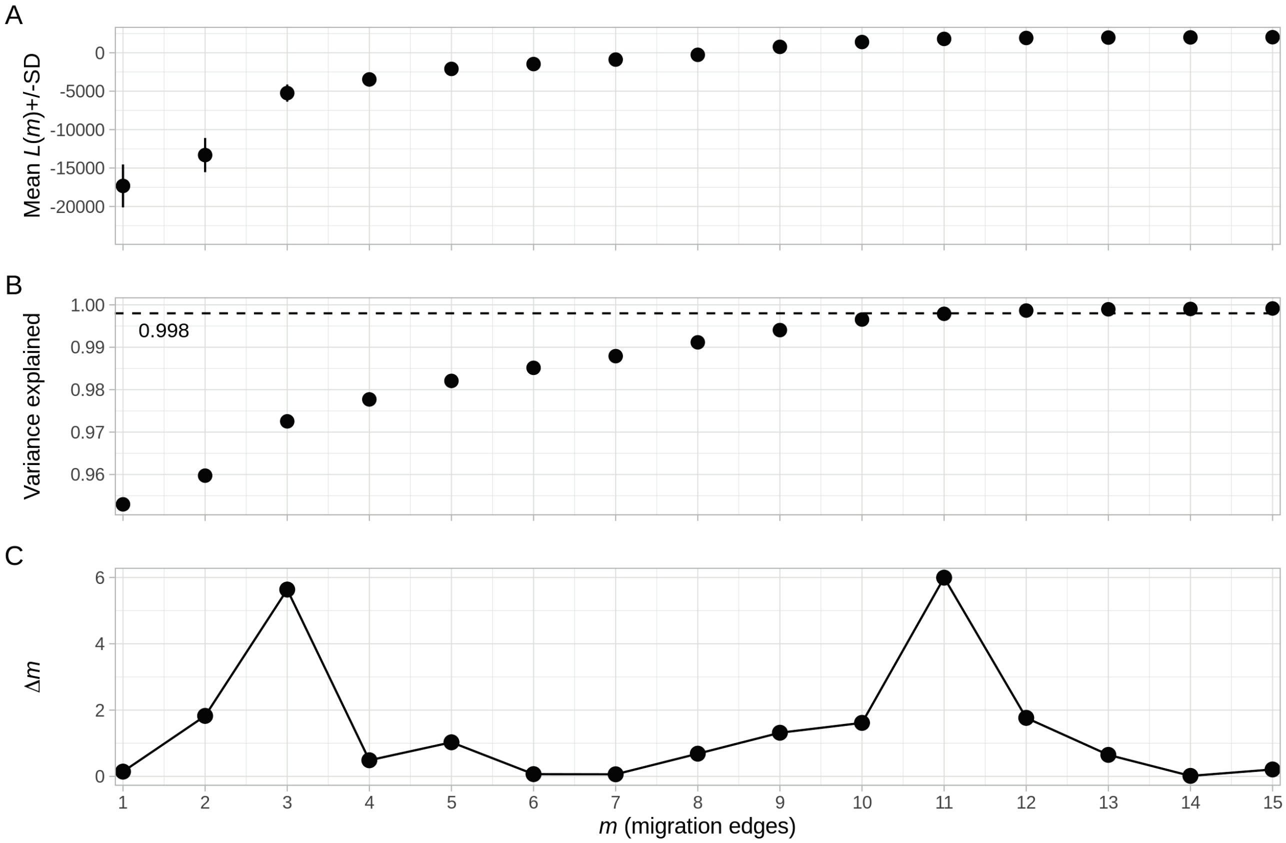


**Figure S10.** OptM results for the low-density SNP data set. **A.** the mean and standard deviation (SD) across 10 iterations for the composite likelihood ($L(m)$). **B.** the proportion of variance explained showing the 99.8% threshold (horizontal dotted line) recommended by Pickrell and Pritchard (2012). **C.** the second-order rate of change ($\Delta m$) across migration edges ($m$).

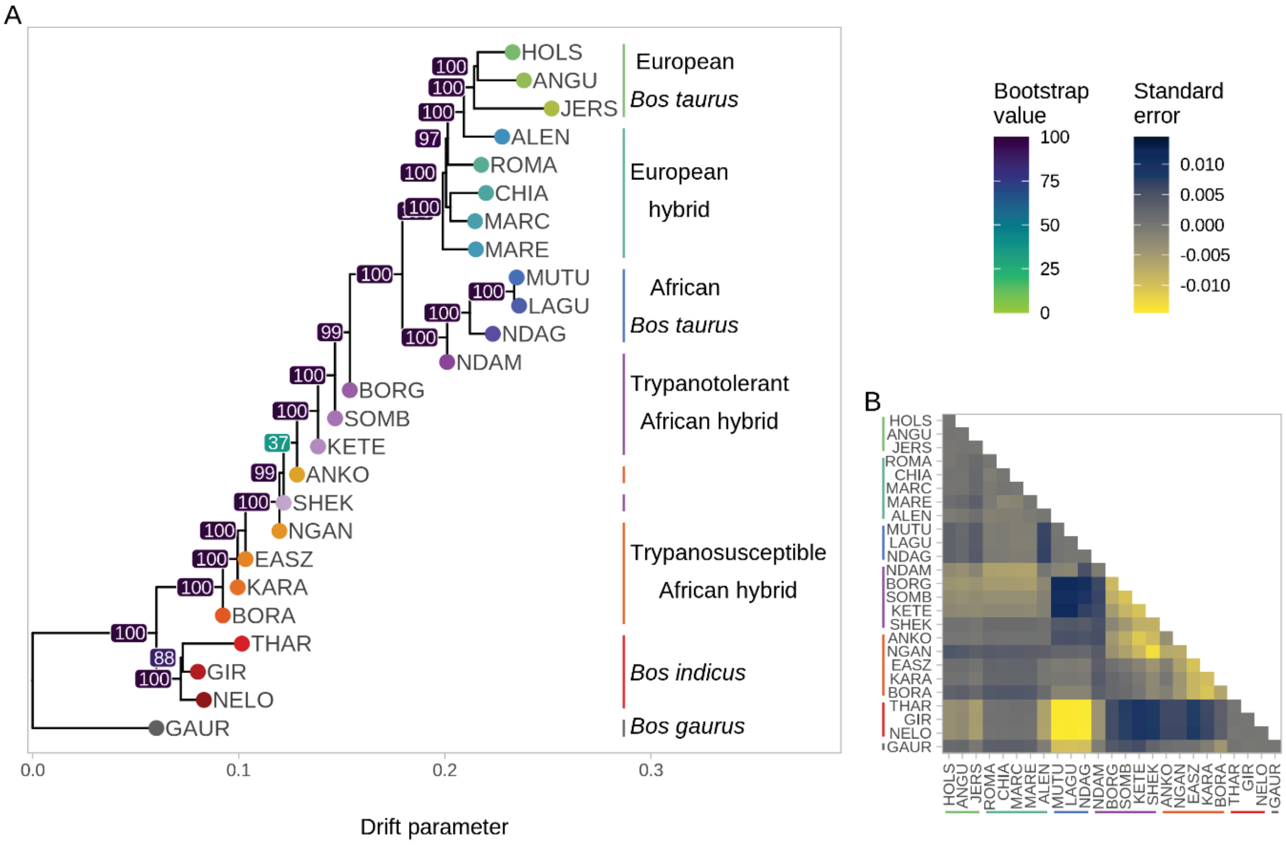


**Figure S11. A.** TreeMix phylogenetic tree for the high-density SNP data set with bootstrap values. **B.** a heatmap showing the standard error.

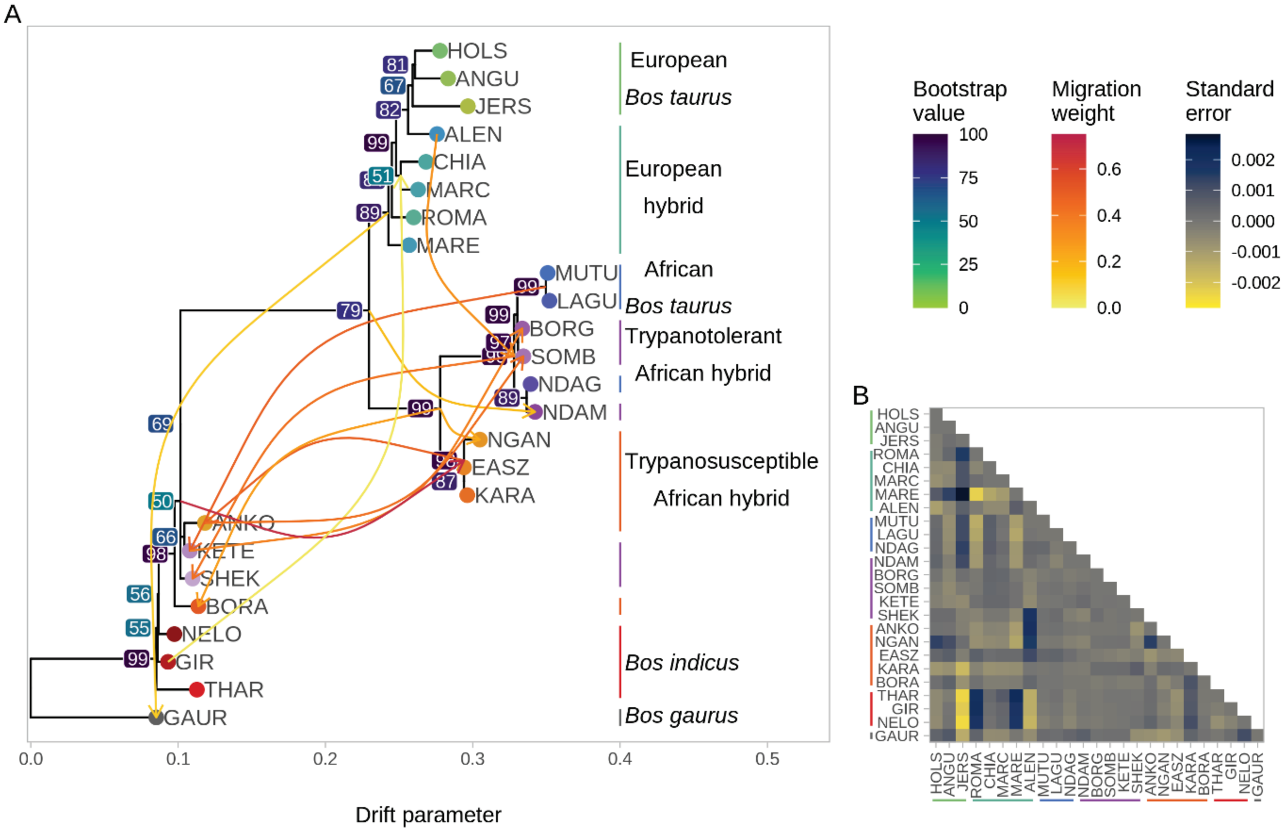


**Figure S12. A.** TreeMix phylogenetic tree for the high-density SNP data set with bootstrap values and 12 migration edges. **B.** a heatmap showing standard error.

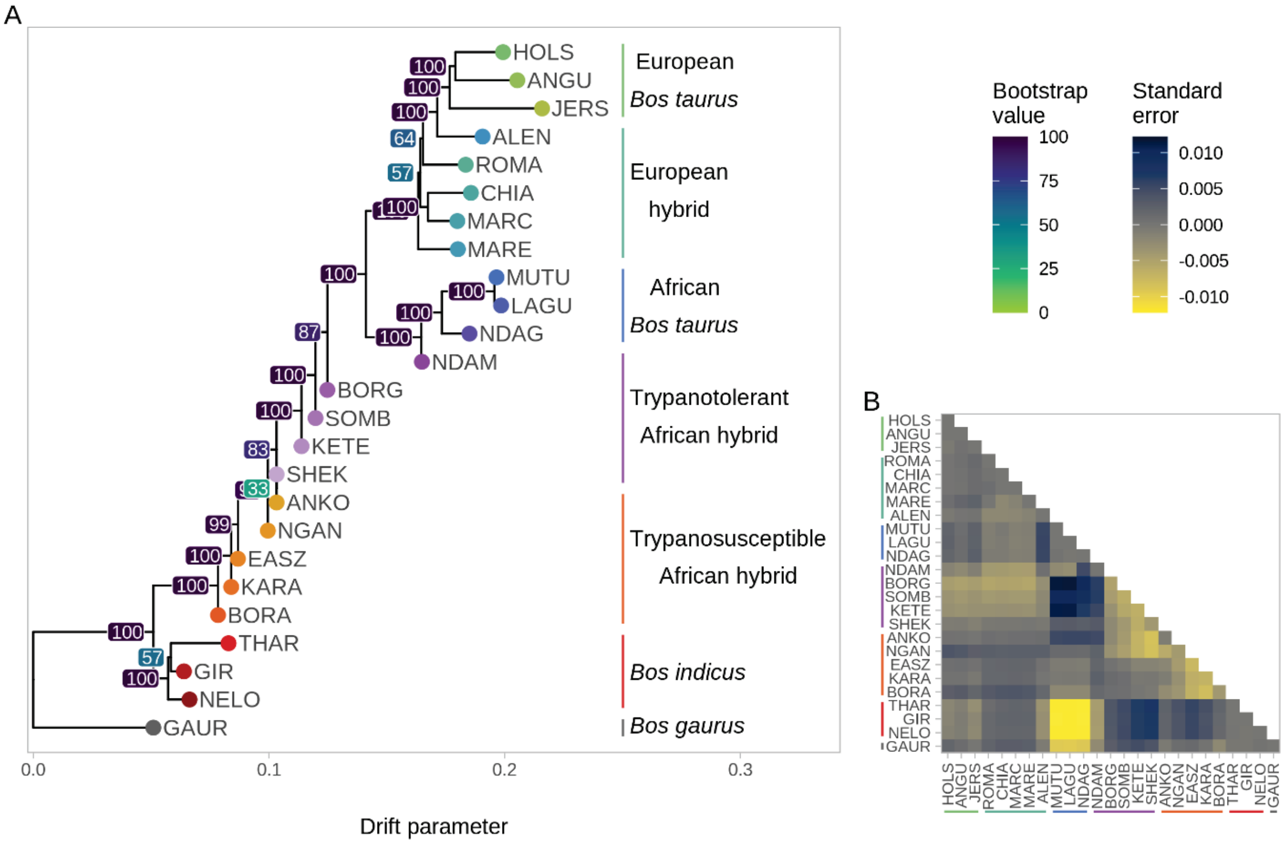


**Figure S13. A.** TreeMix phylogenetic tree for the low-density SNP data set with bootstrap values. **B.** a heatmap showing the standard error.

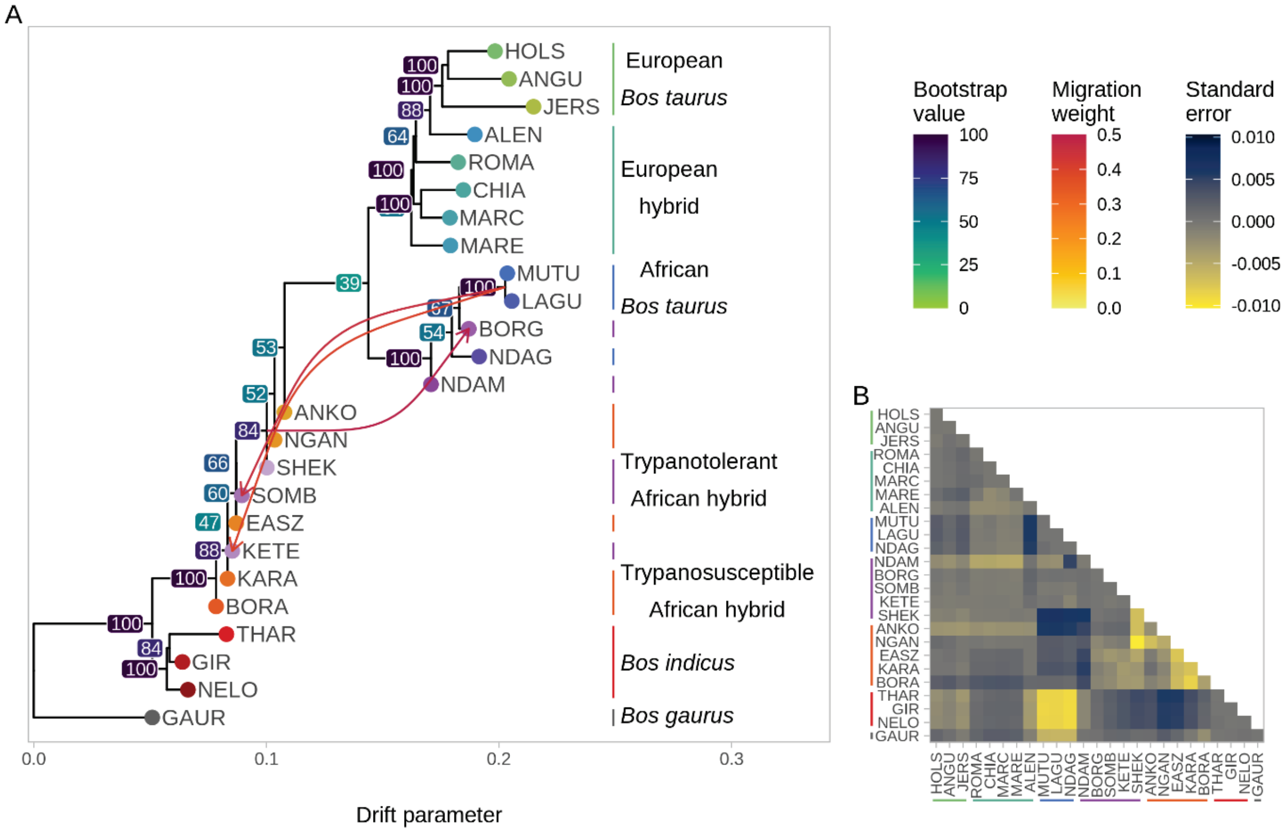


**Figure S14. A.** TreeMix phylogenetic tree for the low-density SNP data set with bootstrap values and 12 migration edges. **B.** a heatmap showing standard error.

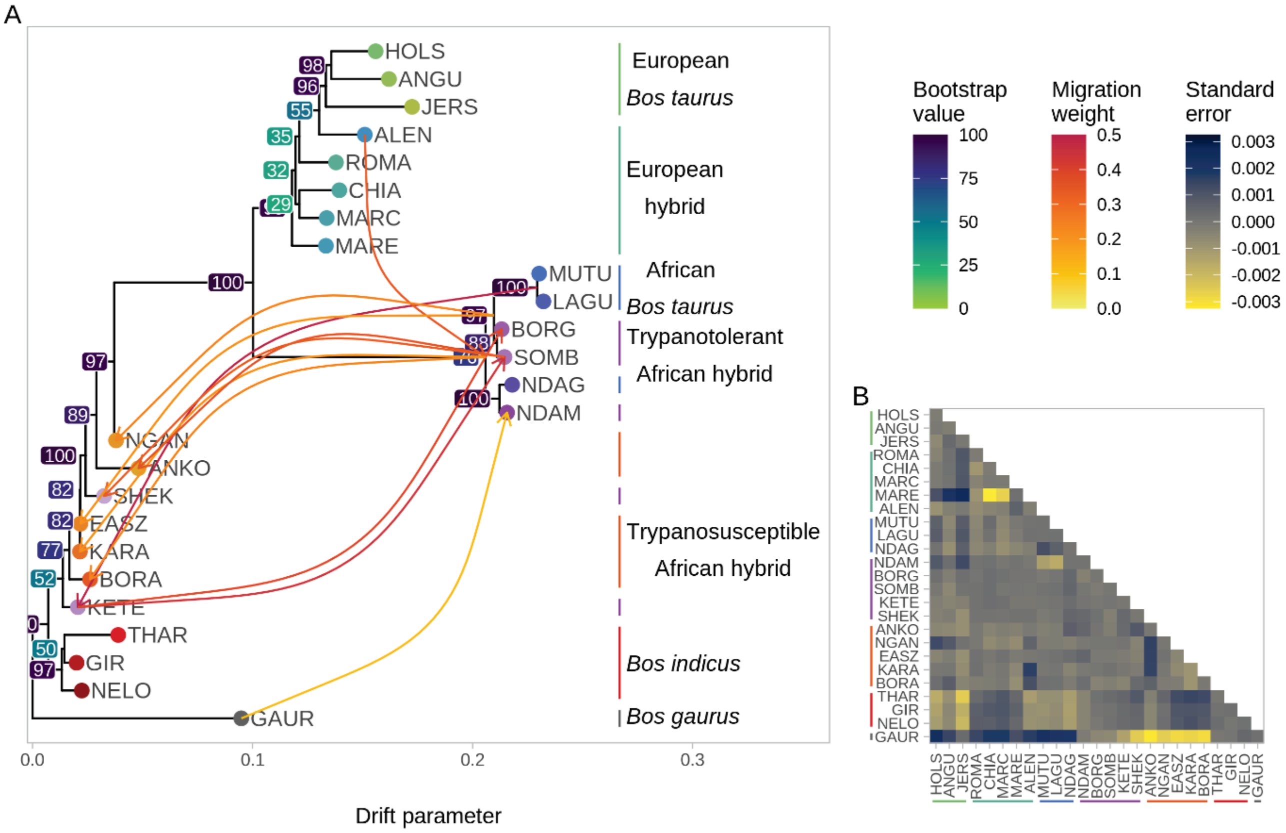


**Figure S15. A.** TreeMix phylogenetic tree for the high-density SNP data set with bootstrap values and 11 migration edges. **B.** a heatmap showing standard error.

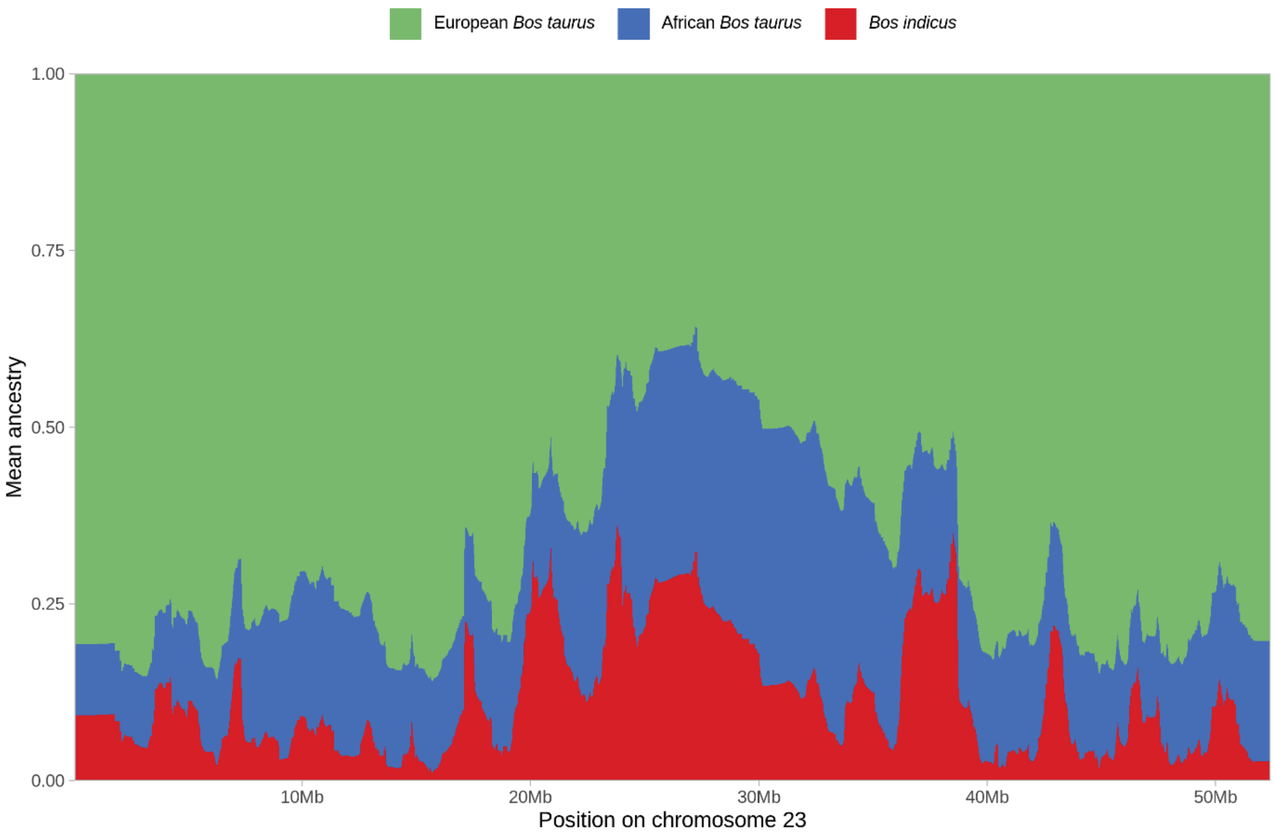


**Figure S16.** Local ancestry results for chromosome 23 (BTA23) for the European hybrid group calculated using MOSAIC with the high-density SNP data set. Each vertical line on the chromosome plot represents a SNP and is coloured according to the ancestry results.

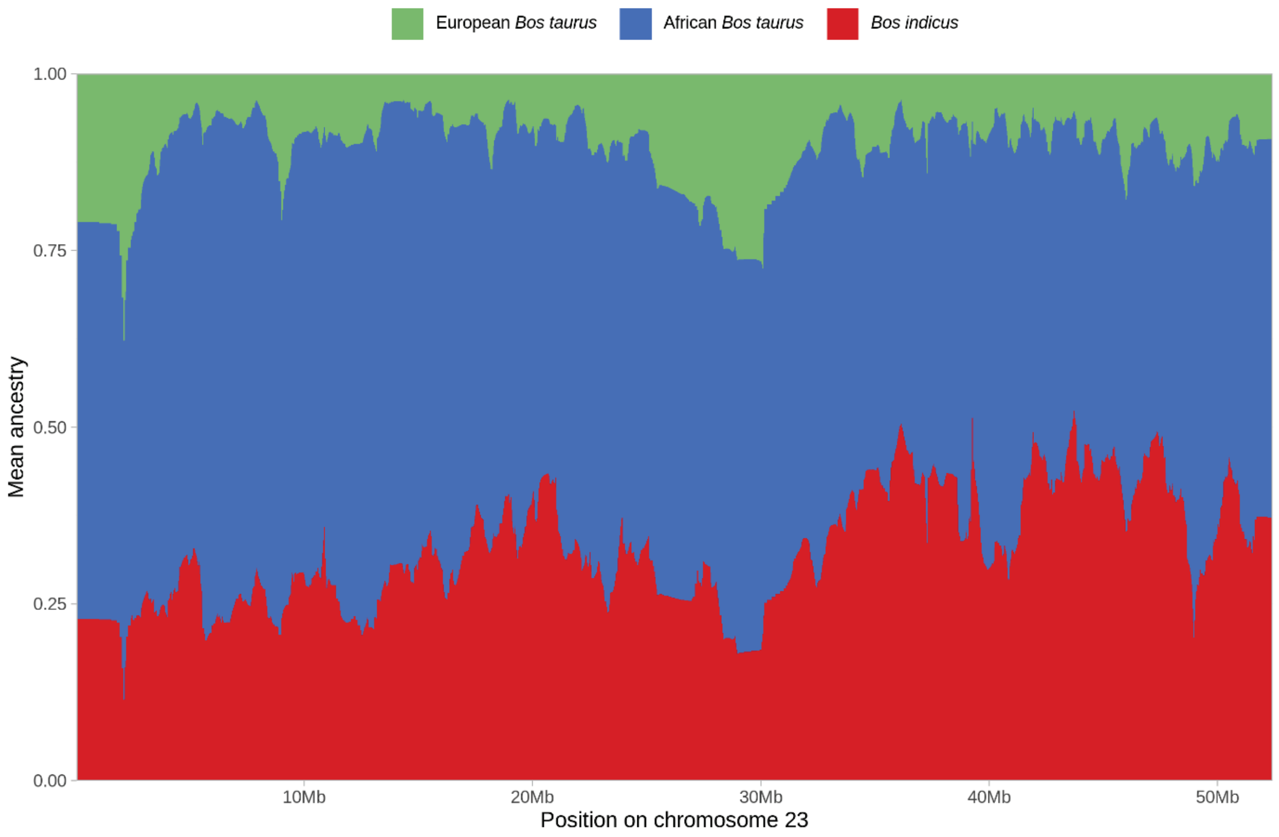


**Figure S17.** Local ancestry results for chromosome 23 (BTA23) for the African trypanotolerant hybrid group calculated using MOSAIC with the high-density SNP data set. Each vertical line on the chromosome plot represents a SNP and is coloured according to the ancestry results.

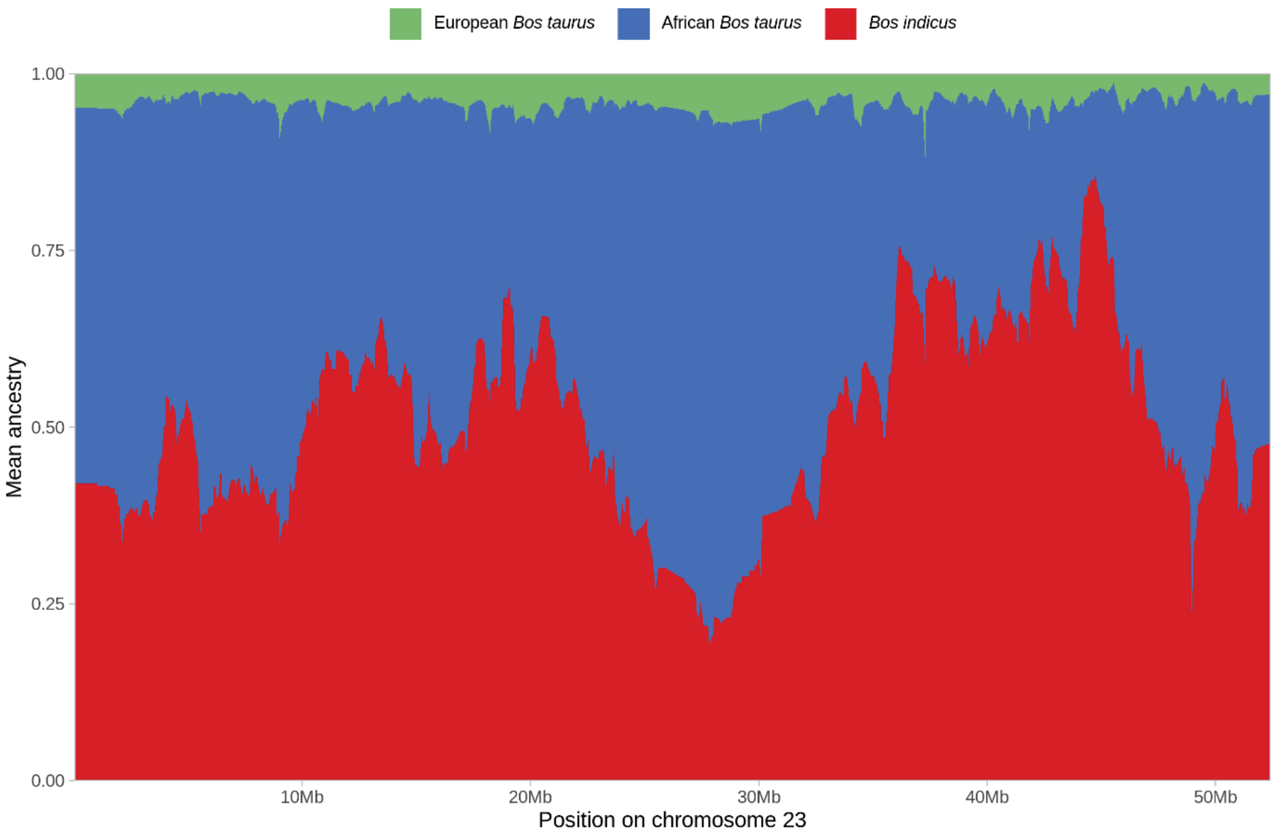


**Figure S18.** Local ancestry results for chromosome 23 (BTA23) for the African trypanosusceptible hybrid group calculated using MOSAIC with the high-density SNP data set. Each vertical line on the chromosome plot represents a SNP and is coloured according to the ancestry results.

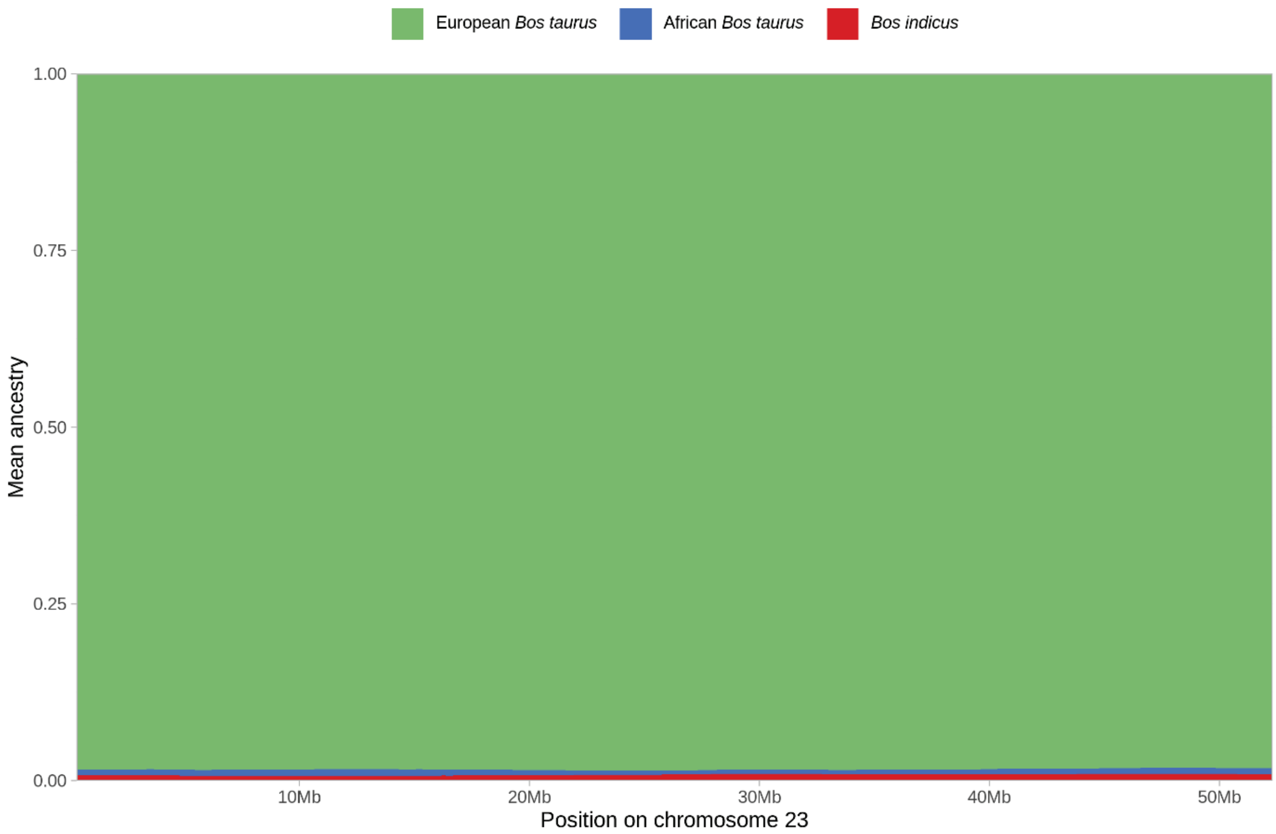


**Figure S19.** Local ancestry results for chromosome 23 (BTA23) for the European hybrid group calculated using MOSAIC with the low-density SNP data set. Each vertical line on the chromosome plot represents a SNP and is coloured according to the ancestry results.

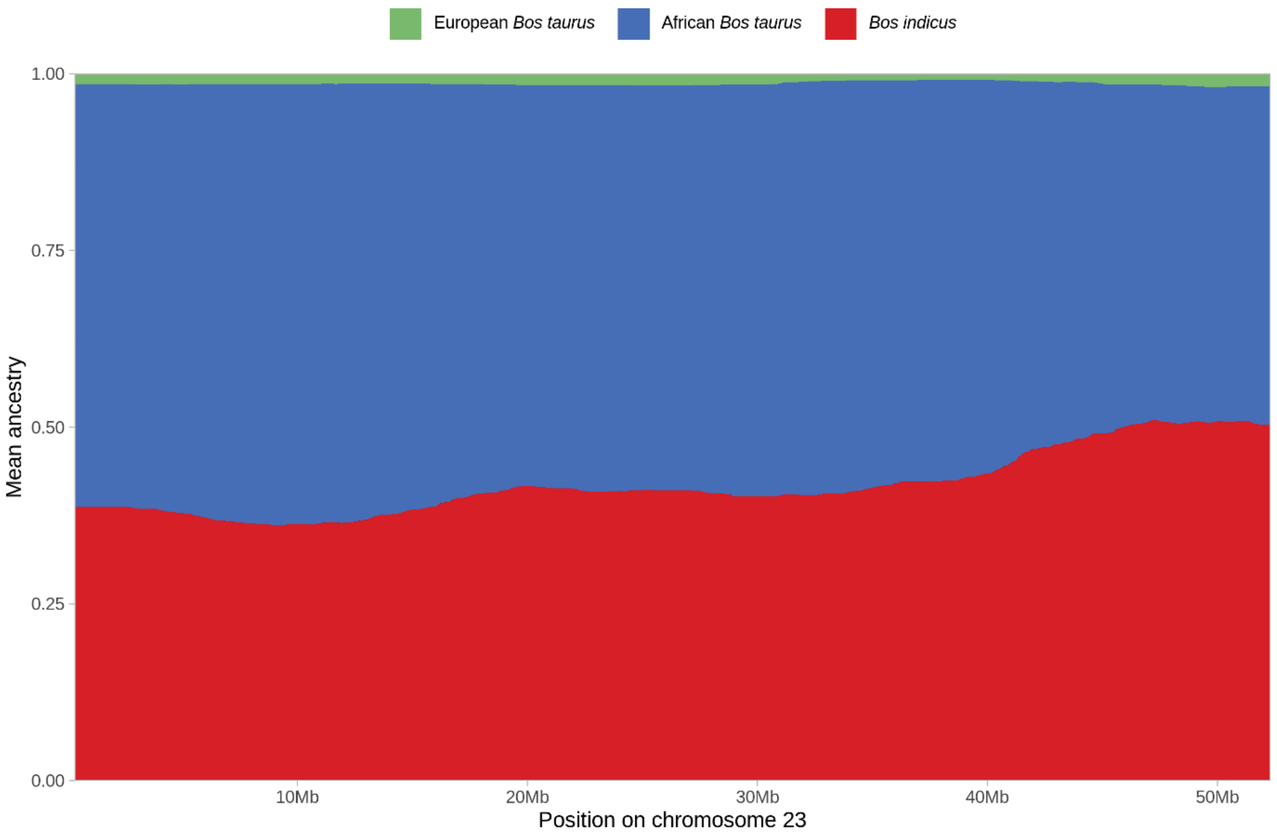


**Figure S20.** Local ancestry results for chromosome 23 (BTA23) for the trypanotolerant African hybrid group calculated using MOSAIC with the low-density SNP data set. Each vertical line on the chromosome plot represents a SNP and is coloured according to the ancestry results.

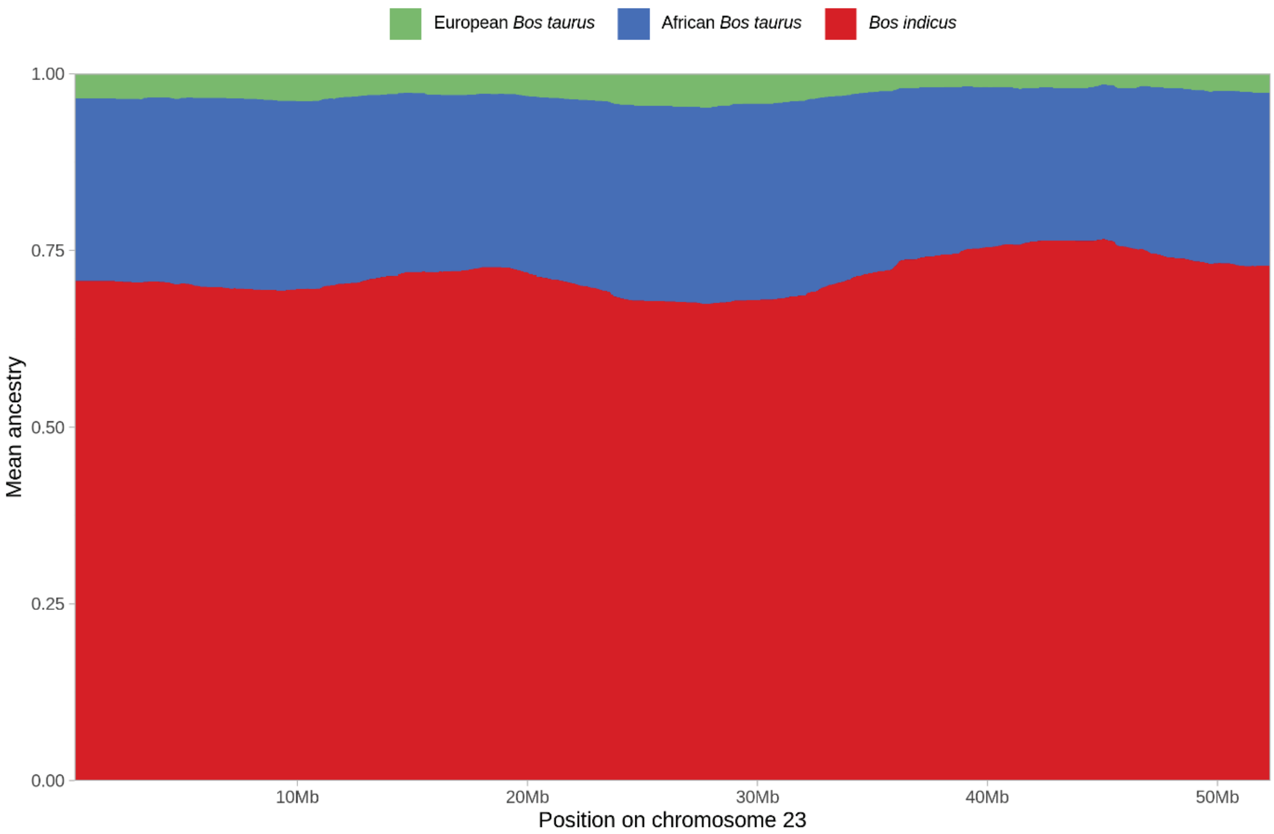


**Figure S21.** Local ancestry results for chromosome 23 (BTA23) for the trypanosusceptible African hybrid group calculated using MOSAIC with the low-density SNP data set. Each vertical line on the chromosome plot represents a SNP and is coloured according to the ancestry results.

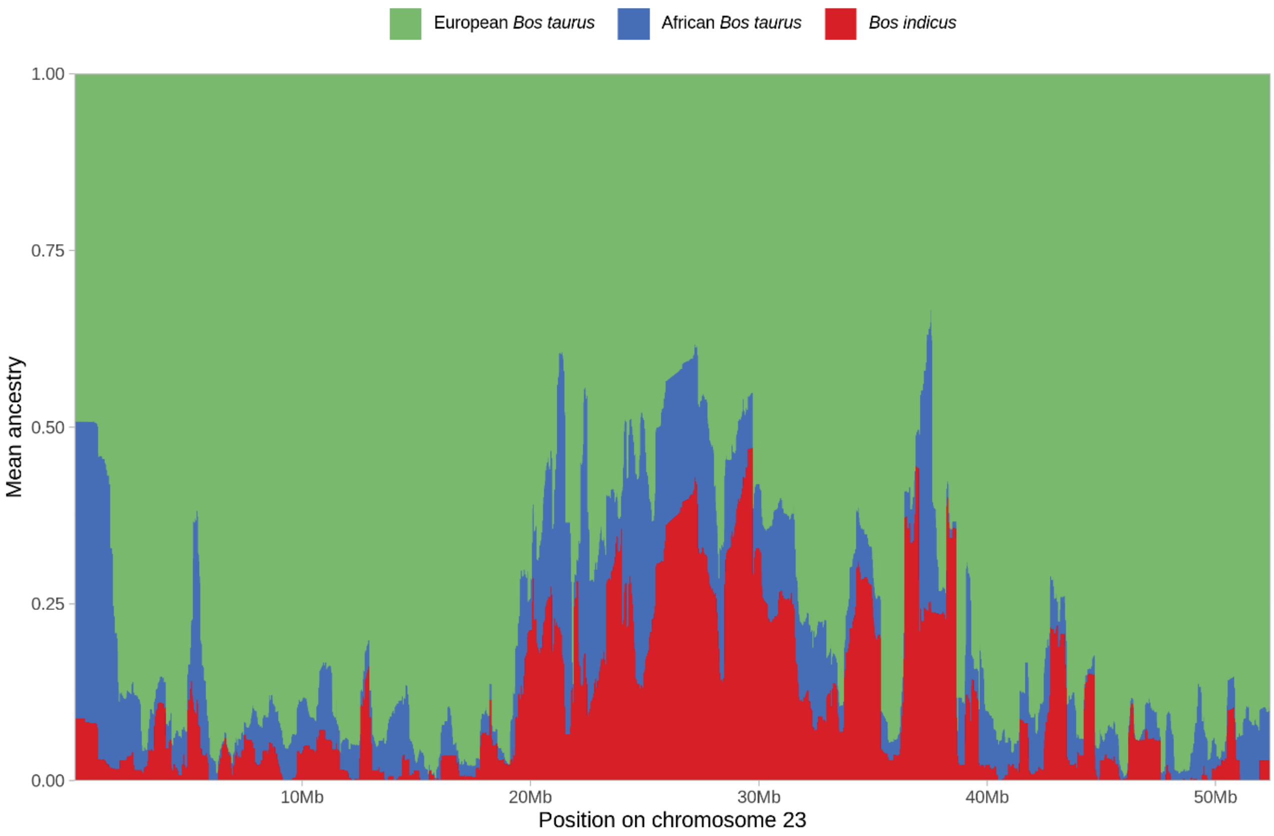


**Figure S22.** Local ancestry results for chromosome 23 (BTA23) for the European hybrid group calculated using ELAI with the high-density SNP data set. Each vertical line on the chromosome plot represents a SNP and is coloured according to the ancestry results.

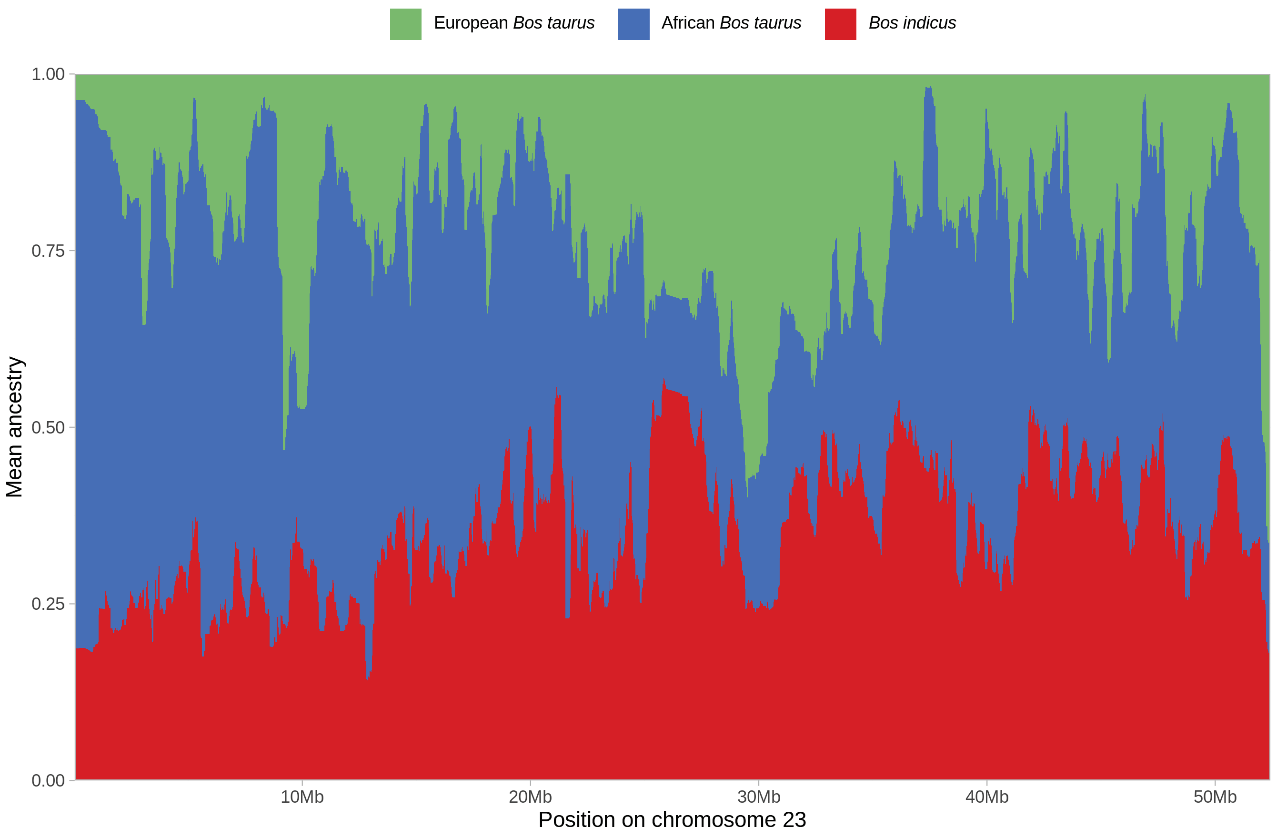


**Figure S23.** Local ancestry results for chromosome 23 (BTA23) for the trypanotolerant African hybrid group calculated using ELAI with the high-density SNP data set. Each vertical line on the chromosome plot represents a SNP and is coloured according to the ancestry results.

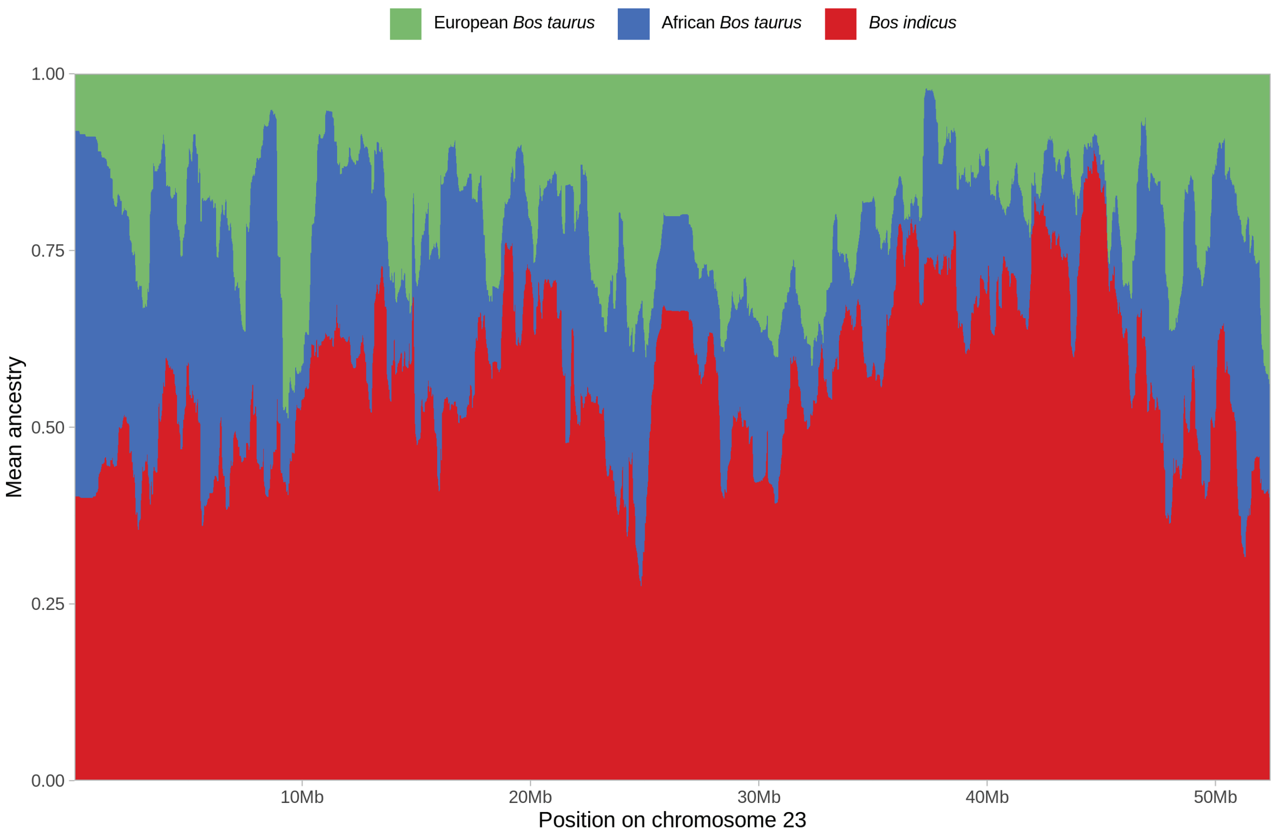


**Figure S24.** Local ancestry results for chromosome 23 (BTA23) for the trypanosusceptible African hybrid group calculated using ELAI with the high-density SNP data set. Each vertical line on the chromosome plot represents a SNP and is coloured according to the ancestry results.

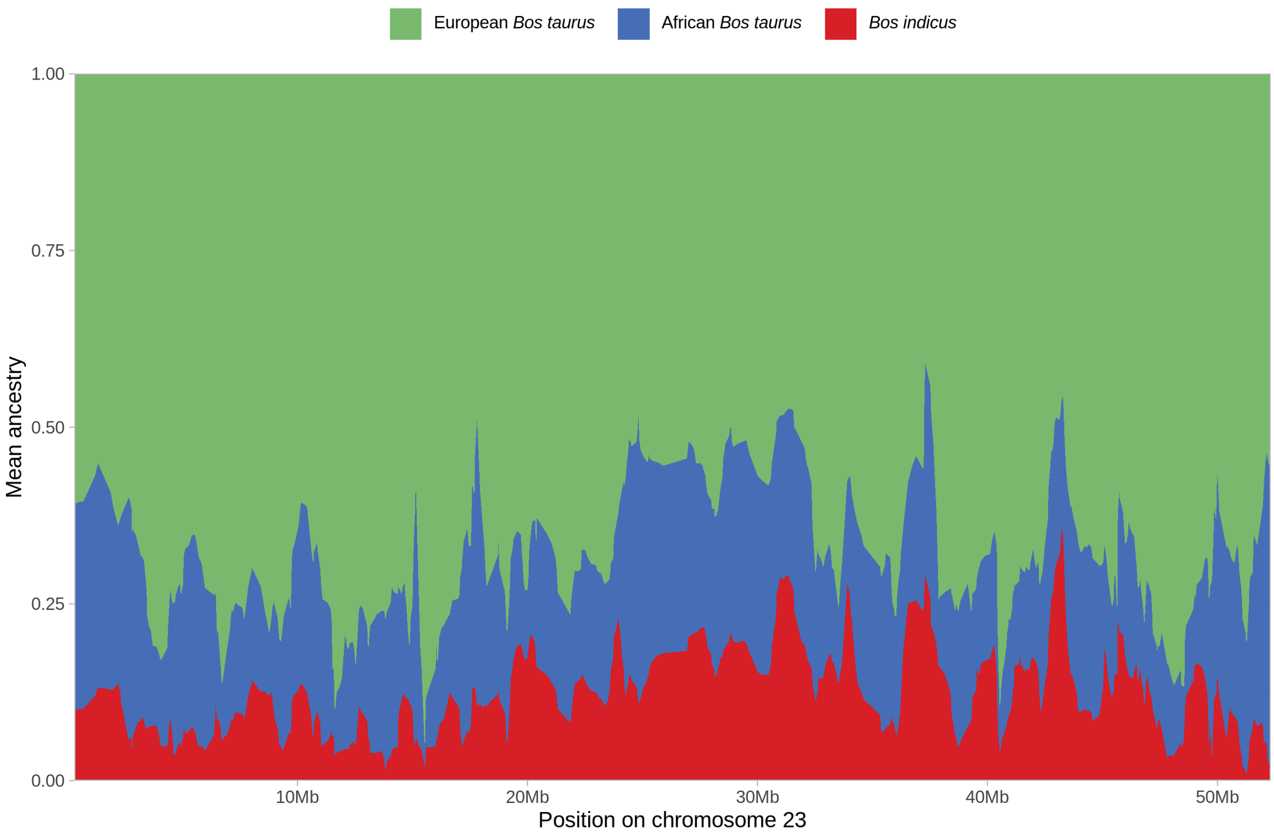


**Figure S25.** Local ancestry results for chromosome 23 (BTA23) for the European hybrid group calculated using ELAI with the low-density SNP data set. Each vertical line on the chromosome plot represents a SNP and is coloured according to the ancestry results.

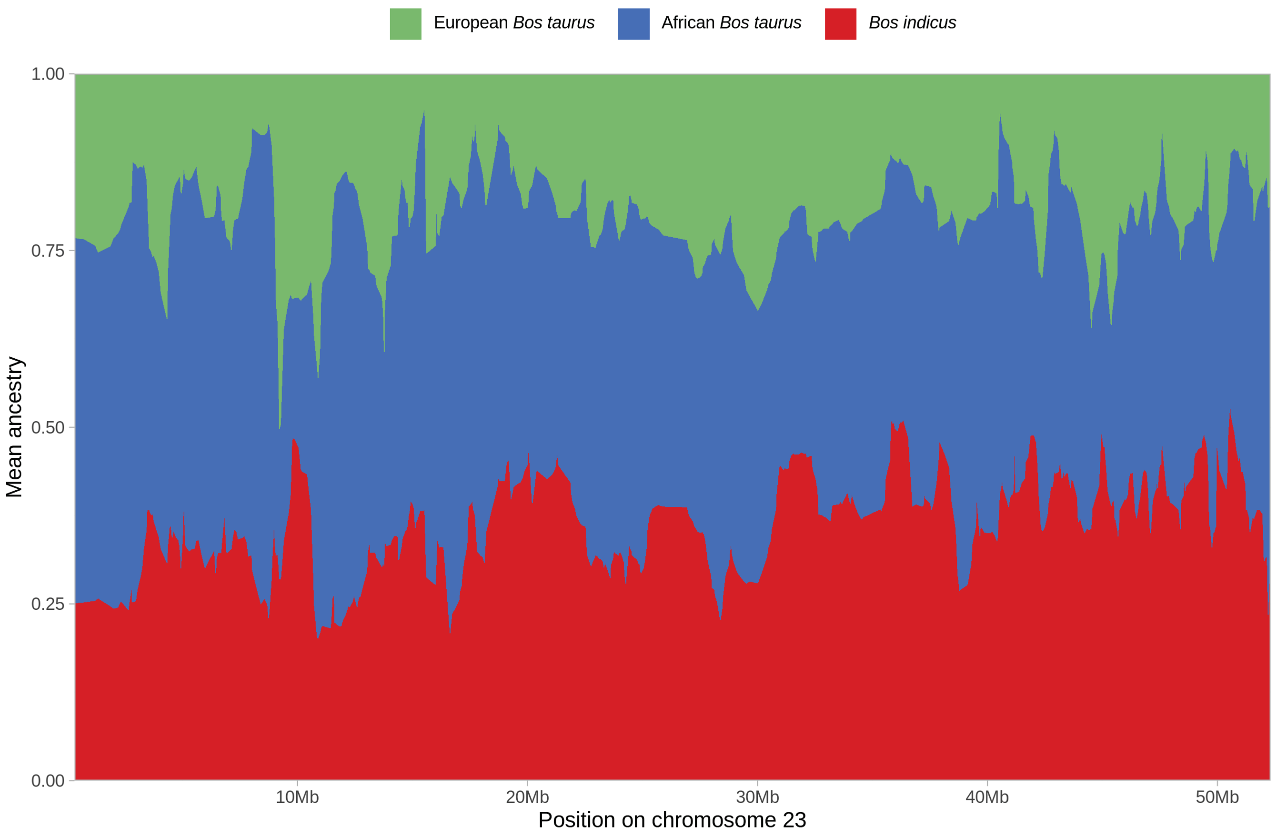


**Figure S26.** Local ancestry results for chromosome 23 (BTA23) for the trypanotolerant African hybrid group calculated using ELAI with the low-density SNP data set. Each vertical line on the chromosome plot represents a SNP and is coloured according to the ancestry results.

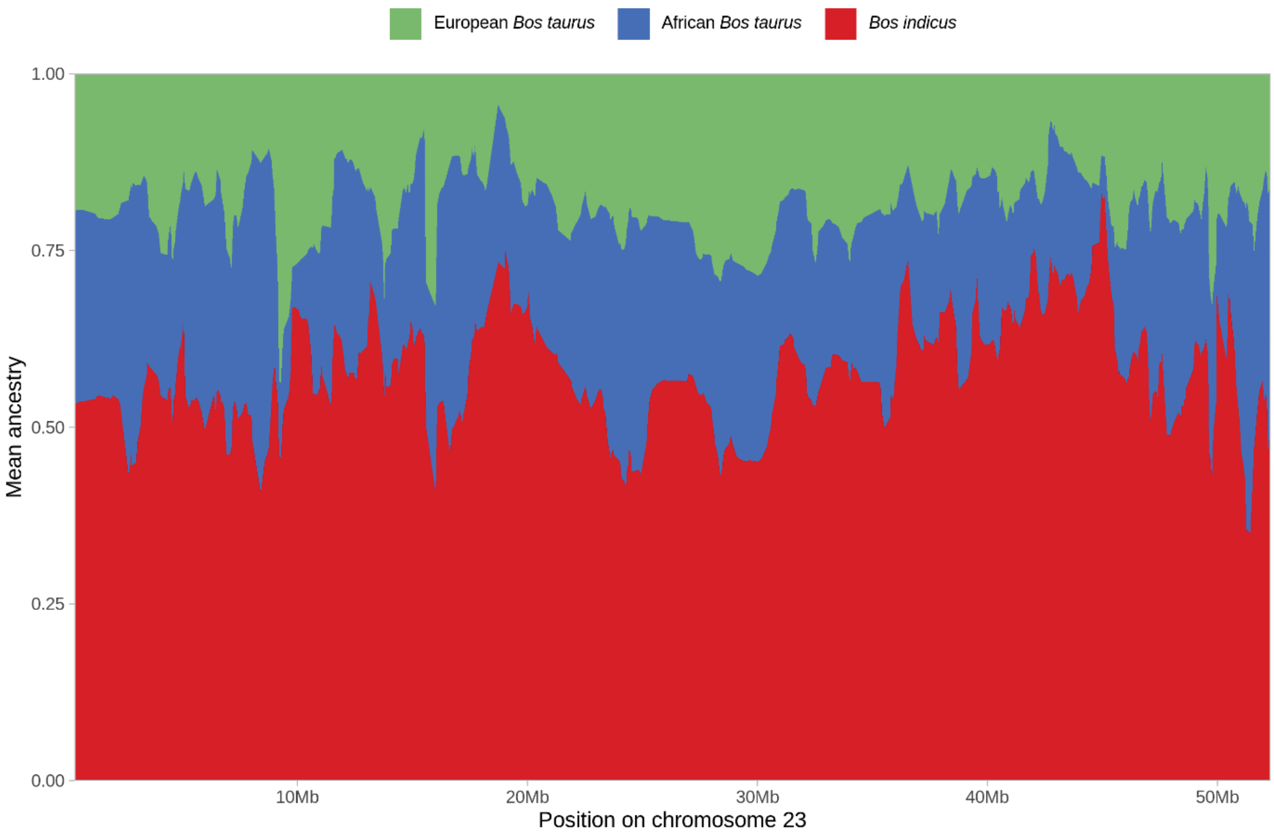


**Figure S27.** Local ancestry results for chromosome 23 (BTA23) for the trypanosusceptible African hybrid group calculated using ELAI with the low-density SNP data set. Each vertical line on the chromosome plot represents a SNP and is coloured according to the ancestry results.

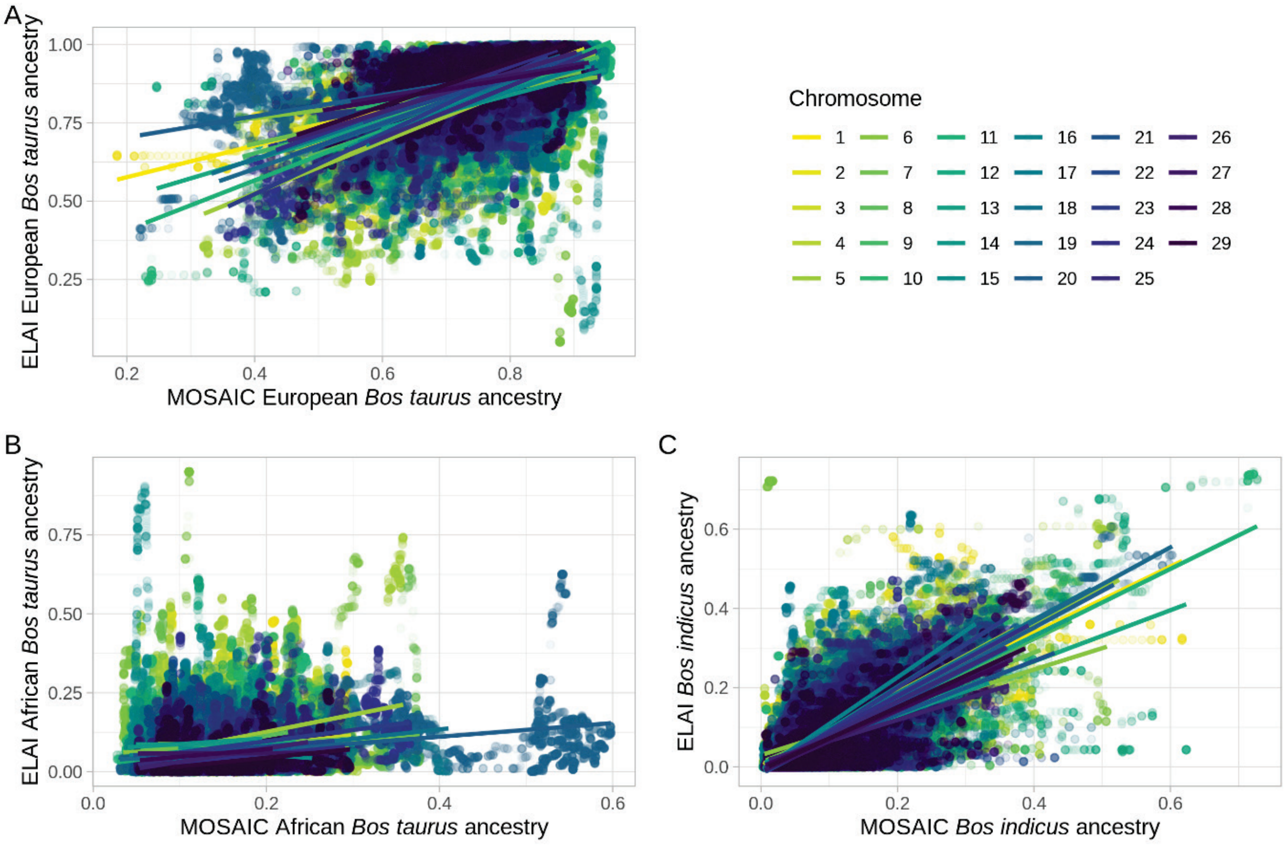


**Figure S28.** Correlation plots for the European hybrid local ancestry results (high-density SNP data set). Each dot represents a SNP coloured according to chromosome. The positions on the x and y-axes indicate the weighted mean ancestry proportions for that SNP according to MOSAIC and ELAI, respectively. **A.** European *Bos taurus*, **B.** African *Bos taurus*, and **C.** *Bos indicus* ancestry components. The lines represent linear models for each chromosome, coloured accordingly.

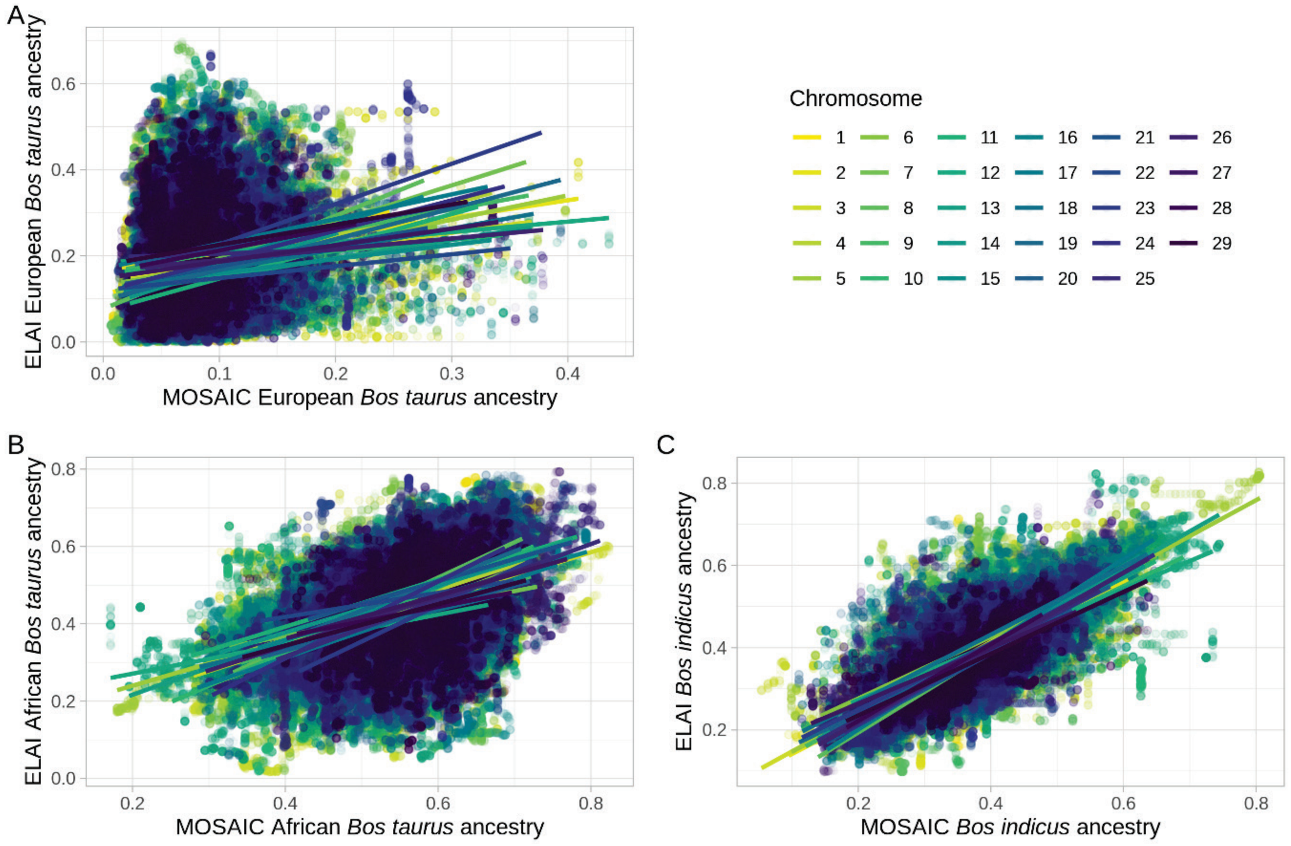


**Figure S29.** Correlation plots for the trypanotolerant African hybrid local ancestry results (high-density SNP data set). Each dot represents a SNP coloured according to chromosome. The positions on the x and y-axes indicate the weighted mean ancestry proportions for that SNP according to MOSAIC and ELAI, respectively. **A.** European *Bos taurus*, **B.** African *Bos taurus*, and **C.** *Bos indicus* ancestry components. The lines represent linear models for each chromosome, coloured accordingly.

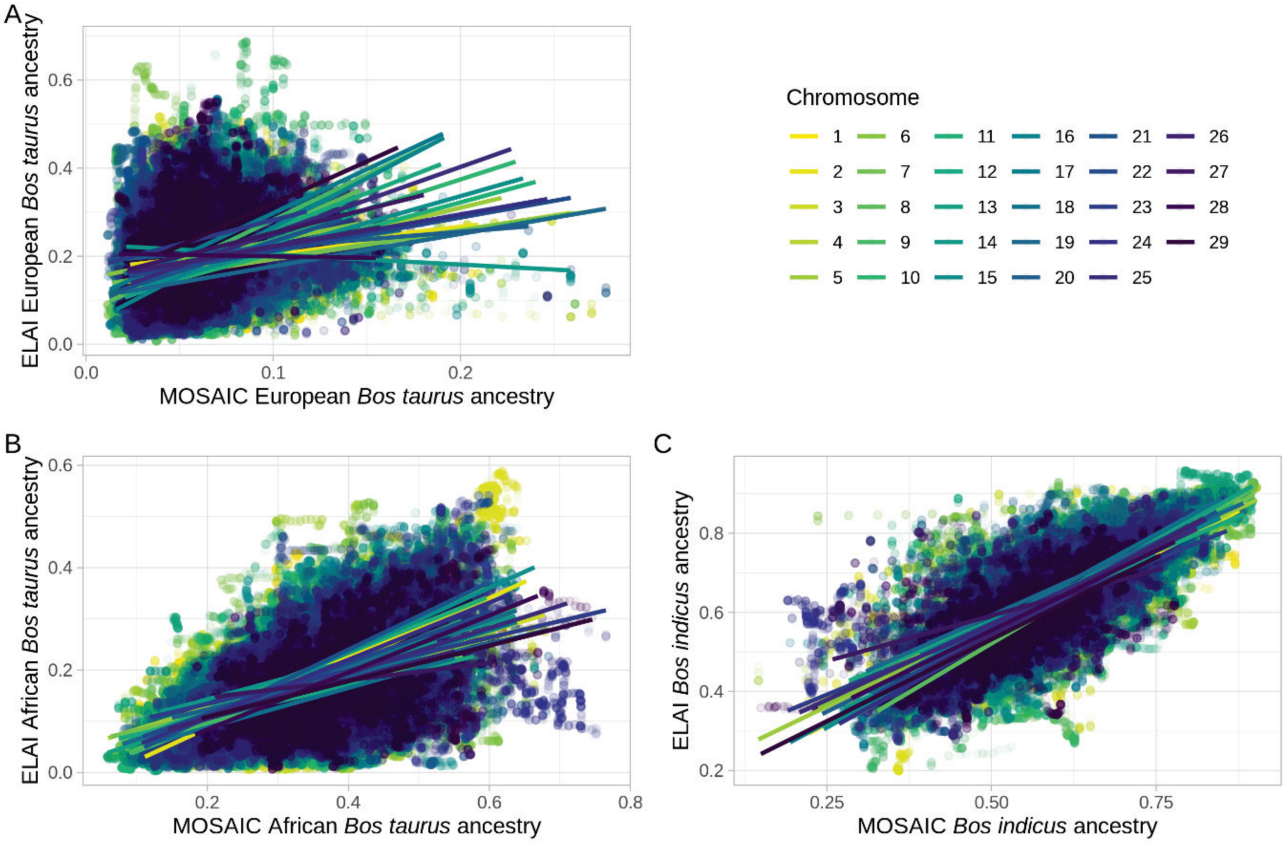


**Figure S30.** Correlation plots for the trypanosusceptible African hybrid local ancestry results (high-density SNP data set). Each dot represents a SNP coloured according to chromosome. The positions on the x and y-axes indicate the weighted mean ancestry proportions for that SNP according to MOSAIC and ELAI, respectively. **A.** European *Bos taurus*, **B.** African *Bos taurus*, and **C.** *Bos indicus* ancestry components. The lines represent linear models for each chromosome, coloured accordingly.

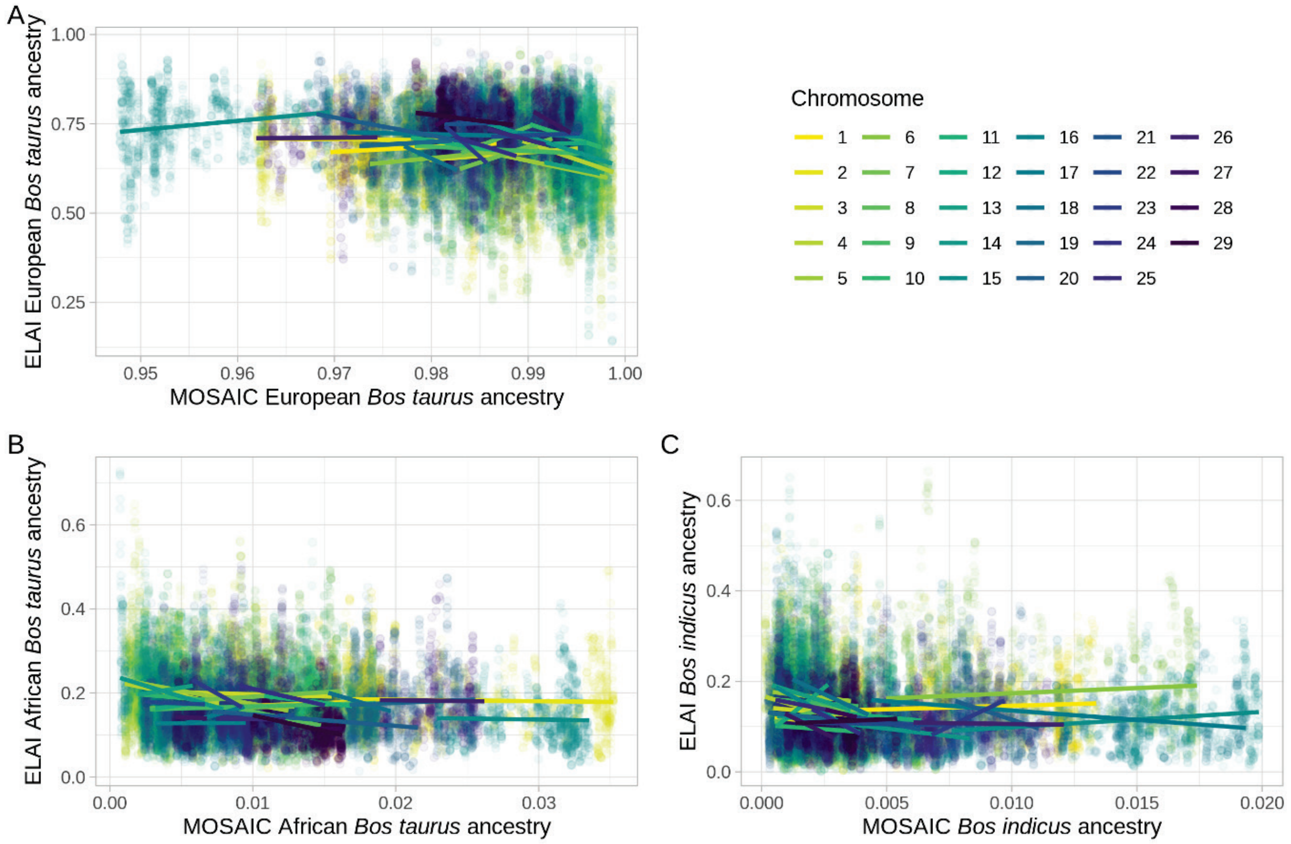


**Figure S31.** Correlation plots for the European hybrid local ancestry results (low-density SNP data set). Each dot represents a SNP coloured according to chromosome. The positions on the x and y-axes indicate the weighted mean ancestry proportions for that SNP according to MOSAIC and ELAI, respectively. **A.** European *Bos taurus*, **B.** African *Bos taurus*, and **C.** *Bos indicus* ancestry components. The lines represent linear models for each chromosome, coloured accordingly.

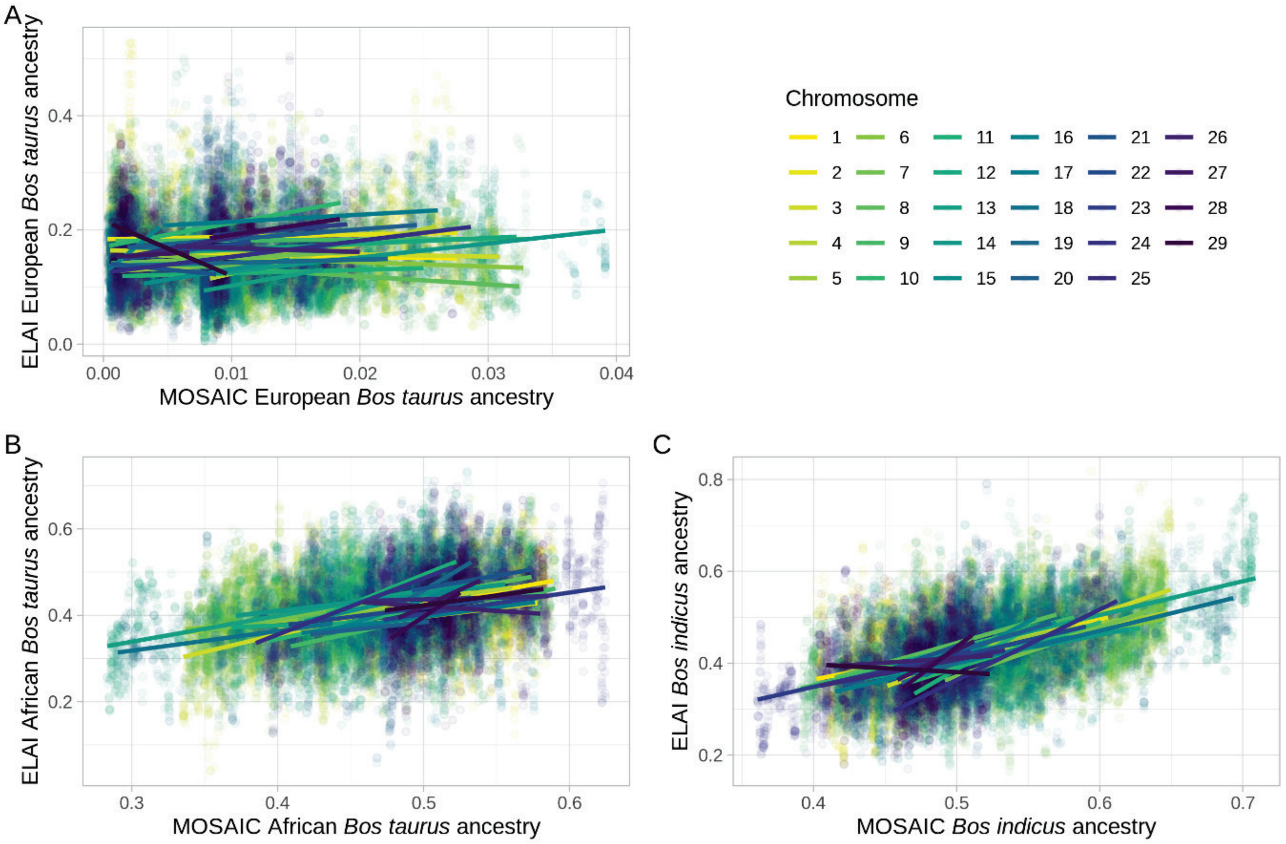


Figure S32. Correlation plots for the trypanotolerant African hybrid local ancestry results (low-density SNP data set). Each dot represents a SNP coloured according to chromosome. The positions on the x and y-axes indicate the weighted mean ancestry proportions for that SNP according to MOSAIC and ELAI, respectively. **A.** European *Bos taurus*, **B.** African *Bos taurus*, and **C.** *Bos indicus* ancestry components. The lines represent linear models for each chromosome, coloured accordingly**.**

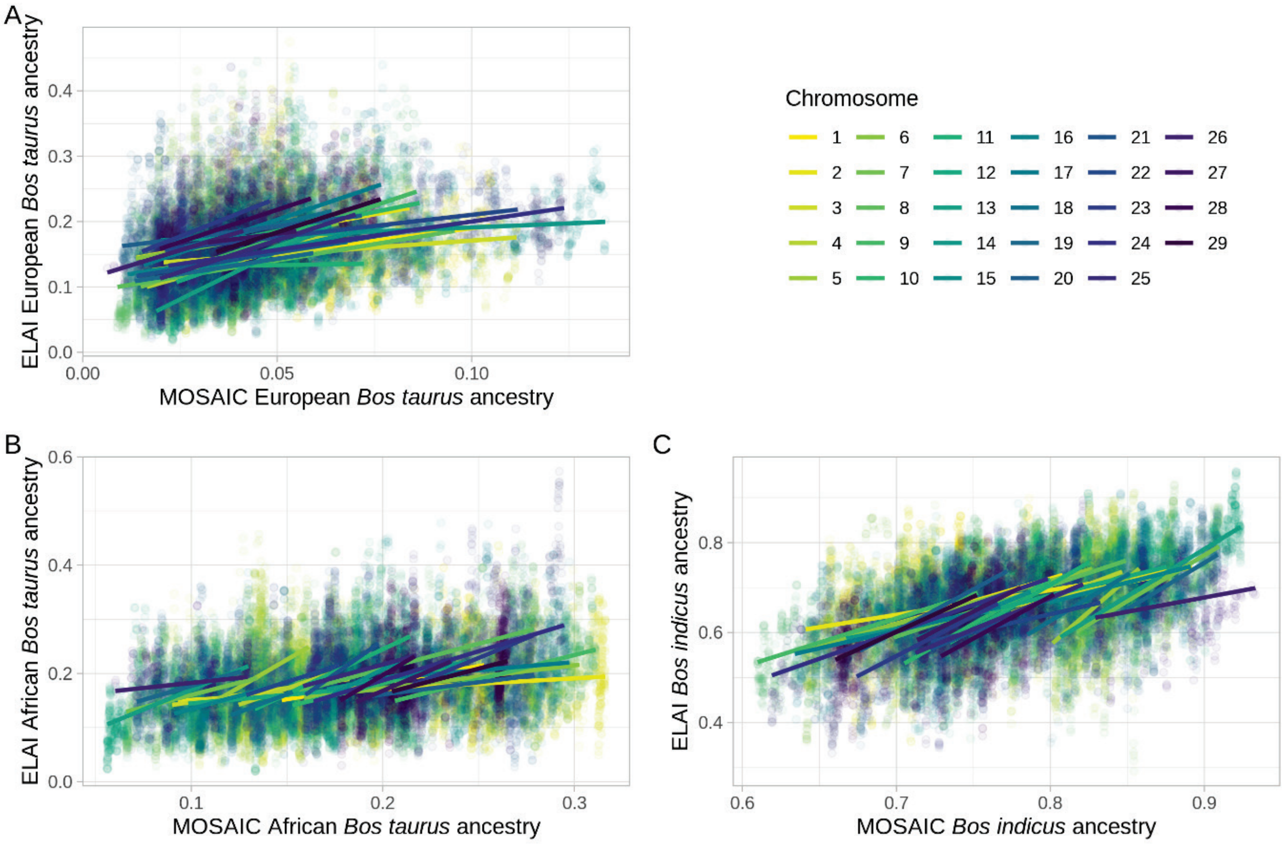


Figure S33. Correlation plots for the trypanosusceptible African hybrid local ancestry results (low-density SNP data set). Each dot represents a SNP coloured according to chromosome. The positions on the x and y-axes indicate the weighted mean ancestry proportions for that SNP according to MOSAIC and ELAI, respectively. **A.** European *Bos taurus*, **B.** African *Bos taurus*, and **C.** *Bos indicus* ancestry components. The lines represent linear models for each chromosome, coloured accordingly**.**

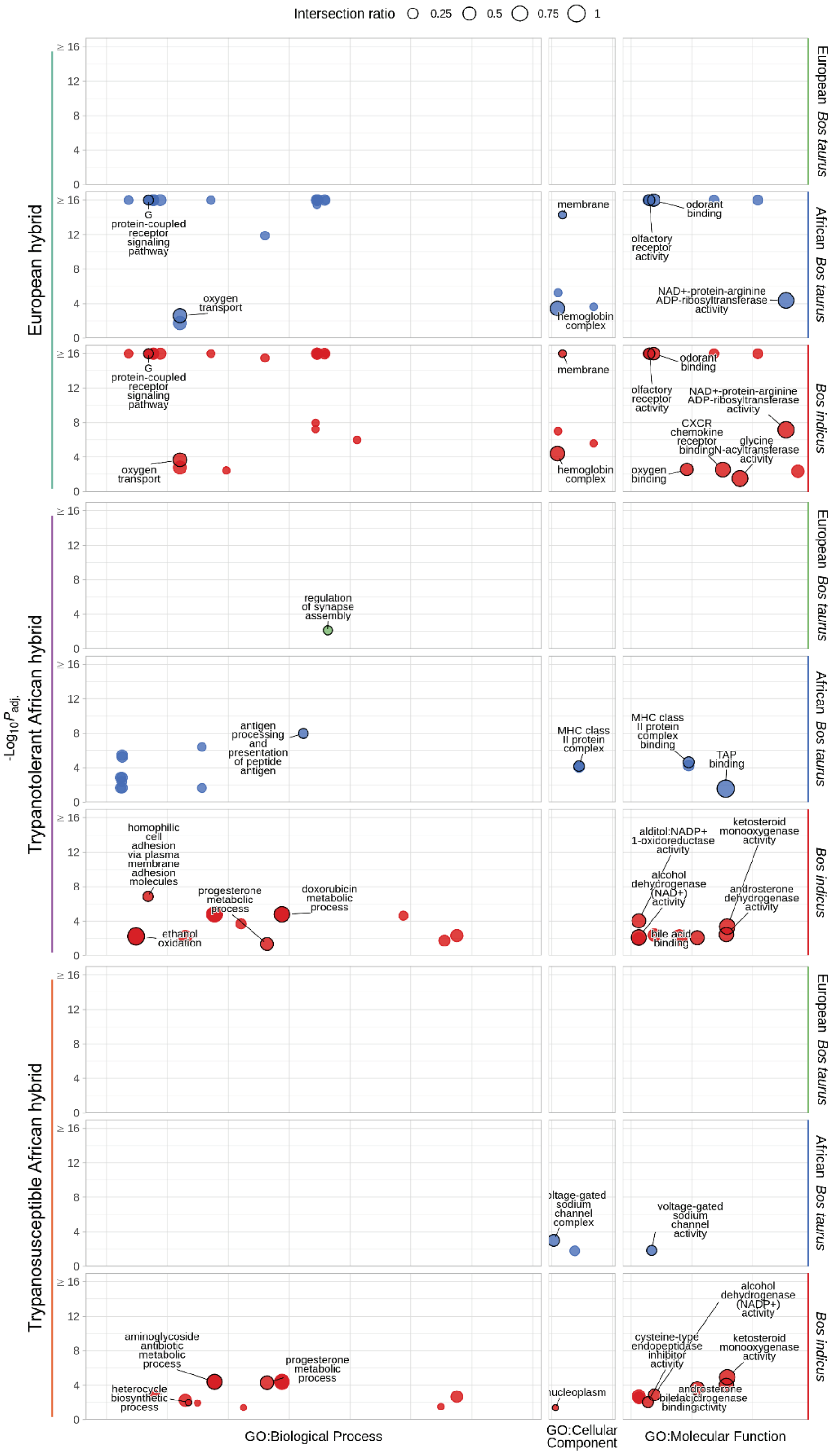


**Figure S34.** g:Profiler functional enrichment of introgressed regions in European, and trypanotolerant and trypanosusceptible African hybrid populations detected with MOSAIC and low-density SNP data. Each circle represents a significantly enriched GO term with the size indicating the ratio of the intersection between the term and the introgressed genes. The y-axis shows the $-\log_{10}(P_{adj.})$ value and the horizontal panels and colours indicate the ancestry component. The vertical panels indicate the source of the term, and position within each panel groups terms from the same GO subtree. The top driver GO terms (up to a maximum of 10) are indicated with a black outline and label.

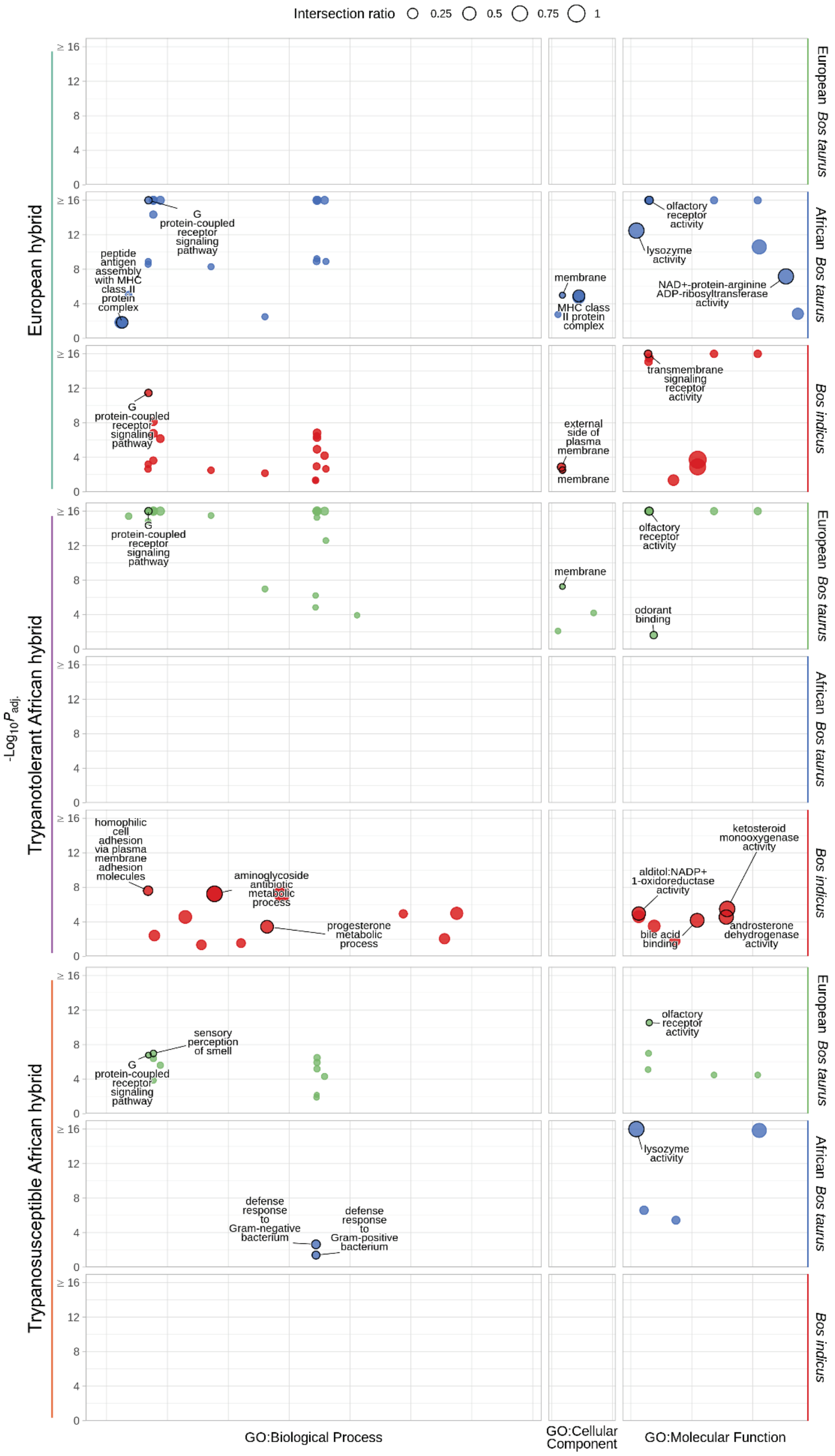


**Figure S35.** g:Profiler functional enrichment of introgressed regions in European, and trypanotolerant and trypanosusceptible African hybrid populations detected with ELAI and high-density SNP data. Each circle represents a significantly enriched GO term with the size indicating the ratio of the intersection between the term and the introgressed genes. The y-axis shows the $-\log_{10}(P_{adj.})$ value and the horizontal panels and colours indicate the ancestry component. The vertical panels indicate the source of the term, and position within each panel groups terms from the same GO subtree. The top driver GO terms (up to a maximum of 10) are indicated with a black outline and label.

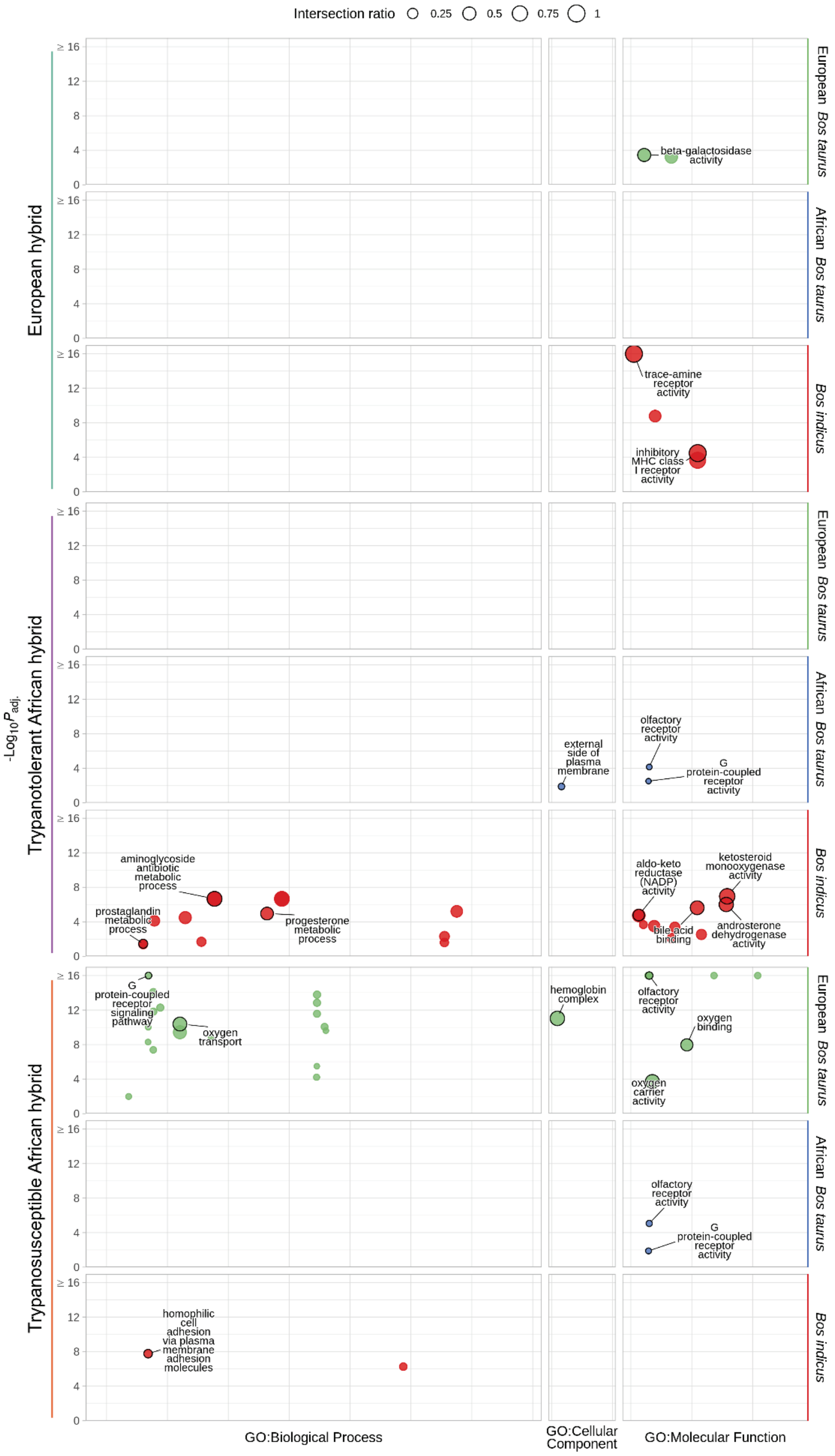


**Figure S36.** g:Profiler functional enrichment of introgressed regions in European, and trypanotolerant and trypanosusceptible African hybrid populations detected with ELAI and low-density SNP data. Each circle represents a significantly enriched GO term with the size indicating the ratio of the intersection between the term and the introgressed genes. The y-axis shows the $-\log_{10}(P_{adj.})$ value and the horizontal panels and colours indicate the ancestry component. The vertical panels indicate the source of the term, and position within each panel groups terms from the same GO subtree. The top driver GO terms (up to a maximum of 10) are indicated with a black outline and label.
